# Multi-omic analysis of guided and unguided forebrain organoids reveal differences in cellular composition and metabolic profiles

**DOI:** 10.1101/2023.12.21.572871

**Authors:** Marie S. Øhlenschlæger, Pia Jensen, Jesper F. Havelund, Magdalena Sutcliffe, Sofie B. Elmkvist, Lucrezia Criscuolo, Steven W. Wingett, Lene A. Jakobsen, Jonathan Brewer, Nils J. Færgeman, Madeline A. Lancaster, Martin R. Larsen, Helle Bogetofte

## Abstract

Neural organoids are invaluable model systems for studying neurodevelopment and neurological diseases. For this purpose, reproducible differentiation protocols are needed that minimize inter-organoid variability whilst generating neural organoids that physiologically resemble the brain area of interest. Currently, two main approaches are used: guided, where the differentiation towards neuroectoderm and subsequently specific CNS regions is driven by applying extrinsic signalling molecules, and unguided, where the intrinsic capability of pluripotent stem cells to generate neuroectoderm without external signalling is promoted. Despite the importance for the field, the resulting differences between these models have not been directly investigated.

To obtain an unbiased comparison, we performed a multi-omic analysis of forebrain organoids generated using a guided and unguided approach focusing on proteomic, lipidomic and metabolomic differences. Furthermore, we characterised differences in phosphorylation and sialylation states of proteins, two key post-translational modifications (PTMs) in neurodevelopment, and performed single cell transcriptomics (scRNAseq). The multi-omic analysis revealed considerable differences in neuronal-, synaptic and glial content, indicating that guided forebrain organoids contain a larger proportion of neurons, including GABAergic interneurons, and synapses whereas unguided organoids contain significantly more GFAP^+^ cells and choroid plexus. Furthermore, substantial differences in mitochondrial- and metabolic profiles were identified, pointing to increased levels of oxidative phosphorylation and fatty acid β-oxidation in unguided forebrain organoids and a higher reliance on glycolysis in guided forebrain organoids.

Overall, our study comprises a thorough description of the multi-omic differences arising when generating guided and unguided forebrain organoids and provide an important resource for the organoid field studying neurodevelopment and -disease.

## Introduction

The development of 3D cell culture techniques has greatly advanced the complexity of *in vitro* models of various tissues. This has markedly benefitted modelling of neuronal tissue, improving the capability to recapitulate key cellular events in early brain development^1–3^. The expanding field of neural organoids was founded by the development of optic cup organoids by the Sasai group in 2011^4^ and was further pioneered by the development of cerebral and forebrain-specific organoids^5–7^. Neural organoids have been used in studies of developmental diseases such as autism spectrum disorder^8,9^, schizophrenia^10^ and microcephaly^5^, and neurodegenerative diseases including Alzheimer’s^11,12^ and Parkinson’s disease^13^. Since the initial development, a large variety of 3D neural organoid protocols have been developed producing an array of models^14–16^ including assembloids, which fuse neural organoids of different regional specifications to simulate connections between regions of the central and peripheral nervous system^17–19^.

Neural organoids can be generated via unguided or guided differentiation. The starting point for both are pluripotent stem cells (PSCs), which as single cells in suspension culture form aggregates, known as embryoid bodies (EBs). The undirected differentiation technique, introduced as cerebral organoids by Lancaster et al. in 2013^5^, relies on the intrinsic ability of PSCs to generate neuroectoderm in the absence of extrinsic signals^5,20^. The EBs are embedded in an extracellular matrix (ECM), which supports the formation of neuroepithelial buds that develop into cortical structures^2,5,20^. The resulting neural organoids can give rise to various brain region identities and contain e.g. retinal tissue and non-neural tissue such as choroid plexus^1,21^. Guided organoid differentiation on the other hand uses small molecules and growth factors to induce regional specification and promote neuronal maturation^6,7^. This enables the formation of brain-region specific organoids resembling dorsal or ventral forebrain, midbrain or hindbrain and is commonly done without ECM embedding^17,22–24^. A commonly used guided protocol is the dorsal forebrain organoid protocol developed by Pasca et al. in 2015^7,17^. The development of commercially available differentiation kits for the dorsal forebrain and cerebral organoid protocols have further facilitated their availability and use. The main differences between this guided and unguided method for generating forebrain organoids (FOs) include the use of dual SMAD inhibition for neural induction, EGF and FGF for neural expansion and later BDNF and NT3 for neuronal maturation in the guided organoid protocol^17^. Furthermore, the unguided protocol includes embedding of EBs in ECM for expansion of large neuroepithelial buds and later orbital shaking of the organoids to increase the flow of nutrients whereas the guided protocol uses static culture in low-attachment plates^5,17^. FOs of both types have been shown to resemble fetal brain tissue on the gene expression level^1,7^, replicate cellular events in cortical plate development and mimic the timing and architecture of early cortical layer formation^5,7,20^. Furthermore, cells of astrocytic lineage have been shown to develop at later timepoints in both types, in line with the timing of *in vivo* brain development^20,25^. The unguided FOs show high diversity with both dorsal and ventral lineages present including the development of GABAergic interneurons, choroid plexus and at later stages also oligodendrocyte precursors^20,21,26^. The guided organoids are expected to develop a more dorsal identity with lower numbers of GABAergic neurons, oligodendrocyte precursors and choroid plexus cells^22,27,3^. Both guided and unguided FOs show spontaneous electrical activity and signs of network formation at later stages of development^5,7,17,21^.

However, our understanding of the differences arising from guided and unguided FO generation is limited as only few studies have included both^28,29^. To establish a more informed foundation for the choice of model system, we therefore performed a direct comparison of the two widely used approaches using a multi-omics methodology including both proteomics, metabolomics, lipidomics and single-cell transcriptomics. Our large-scale proteomic analysis included quantification of two types of post-translational modifications (PTMs), protein phosphorylation, a key indicator of intracellular signalling, and sialylated N-linked glycosylation, which is highly important for neural development and physiology^30^. The comparison was performed early in the differentiation (day 40 to day 120) intending to detect early signs of changes in trajectories between the two approaches. The analyses collectively showed increased neuronal differentiation in the guided FOs and enhanced glial (GFAP^+^) content in the unguided FOs. Unguided FOs contained a substantially larger proportion of cortical hem and the cell populations induced by its signalling in the form of choroid plexus and Cajal Retzius neurons^31^. Surprisingly, the presence of GABAergic interneurons was considerably higher in guided FOs than unguided. The analysis furthermore revealed distinct metabolic activity between the two organoid types with higher levels of oxidative phosphorylation (OXPHOS) and fatty acid β-oxidation (FAO) in unguided FOs and increased glycolysis/penthose phosphate pathway (PPP) reliance in guided FOs.

These results highlight some key differences between guided and unguided FOs and demonstrate the need for in-depth characterization and comparison of organoid models to enable informed decisions on their applications.

## Results

### Guided and unguided FOs differ in size and macroscopic architecture

To examine the differences resulting from generating FOs using the guided or unguided approach, we differentiated FOs of both types in parallel starting from the same iPSCs using commercially available kits from Stem Cell Technologies (Fig. S1A). The expression levels of key neurodevelopmental markers were overall comparable in guided and unguided FOs at day 40 and day 80 as examined by immunocytochemistry (ICC) (Fig.1). Apical-basal polarity was evident with neural cadherin (ncad) expressed at the apical surface of neural rosettes in both types of FOs, and in more ventricular structures seen mainly in the unguided FOs (Fig. 1A), and basal expression of the neuronal markers doublecortin (Dcx), microtubule associated protein 2 (Map2) and NeuN (Fig. 1A-C). The cortical plate marker Tbr1 was expressed by immature neurons at day 40 (Fig. 1C) and the cortical layer markers Satb2 and Ctip2 by separate neuronal populations at day 80 (Fig. 1D). Despite starting from a lower number of iPSCs per organoid, the unguided FOs were significantly larger than the guided ones at day 40 (Fig. S1A-C). Macroscopically, the guided FOs appeared to consist mainly of neural rosettes with adjacent newly-formed neurons, whereas the unguided FOs were less uniform with areas with larger ventricular structures and others with neurons (Fig. 1A-C).

**Figure 1:**
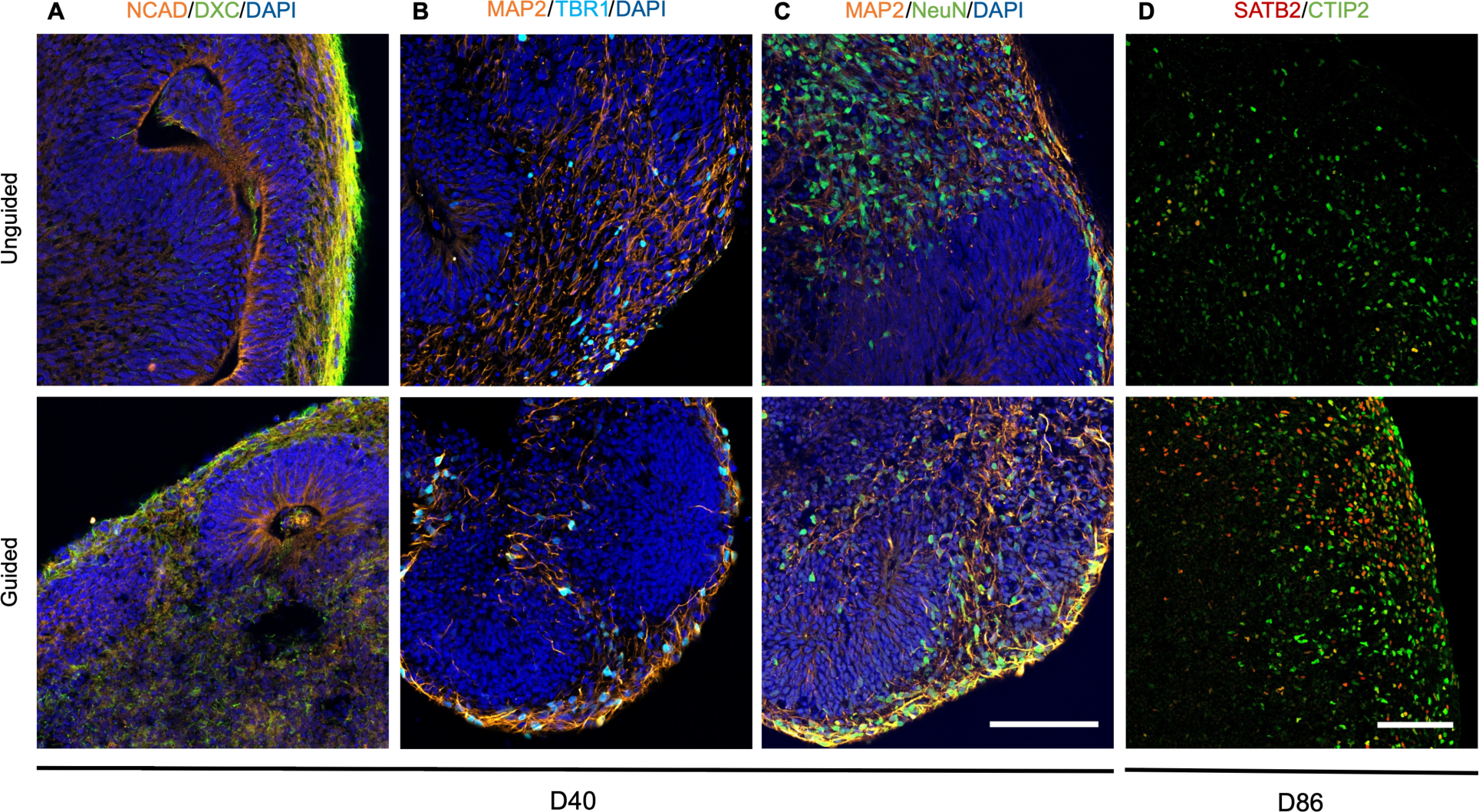
Markers of neuronal maturity and cortical development are expressed in guided and unguided forebrain organoids (FOs) (A-D) Immunocytochemistry of guided and unguided FOs for (A) N-cadherin (Ncad, orange) and doublecortin (Dxc, green), (B) microtubule associated protein 2 (Map2, orange) and T-Box Brain Transcription Factor 1 (Tbr1, light blue), (C) Map2 (orange) and neuronal nuclear protein (NeuN, green), and (D) DNA-binding protein Satb2 (orange) and Ctip2 (green) at differentiation (A-C) day 40 with DAPI (dark blue) and (D) day 80. Scalebars = 100μm.

### Proteomic analysis identifies differences in expression of neuronal and metabolic proteins

With the aim of performing an unbiased analysis of differences between guided and unguided FOs, we subjected day 40 FOs from three independent differentiations to large-scale proteomic analysis (n=9). Using our previously published PTMomics method^32^, we quantified levels of 7082 proteins, 15775 phospho-peptides (from 4238 proteins) and 663 sialylated N-linked glyco-peptides (from 441 proteins) (Fig. 2A-C). Principal component analysis (PCA) clearly separated the unguided and guided FOs based on the non-modified proteins and, although less concisely, also the PTM peptides (Fig. 2D-F). Levels of 757 proteins, 1079 phospho-peptides and 44 sialylated peptides were significantly different between the guided and unguided FOs generated from the same starting iPSCs, highlighting that the differentiation approach can clearly affect the outcome.

**Figure 2:**
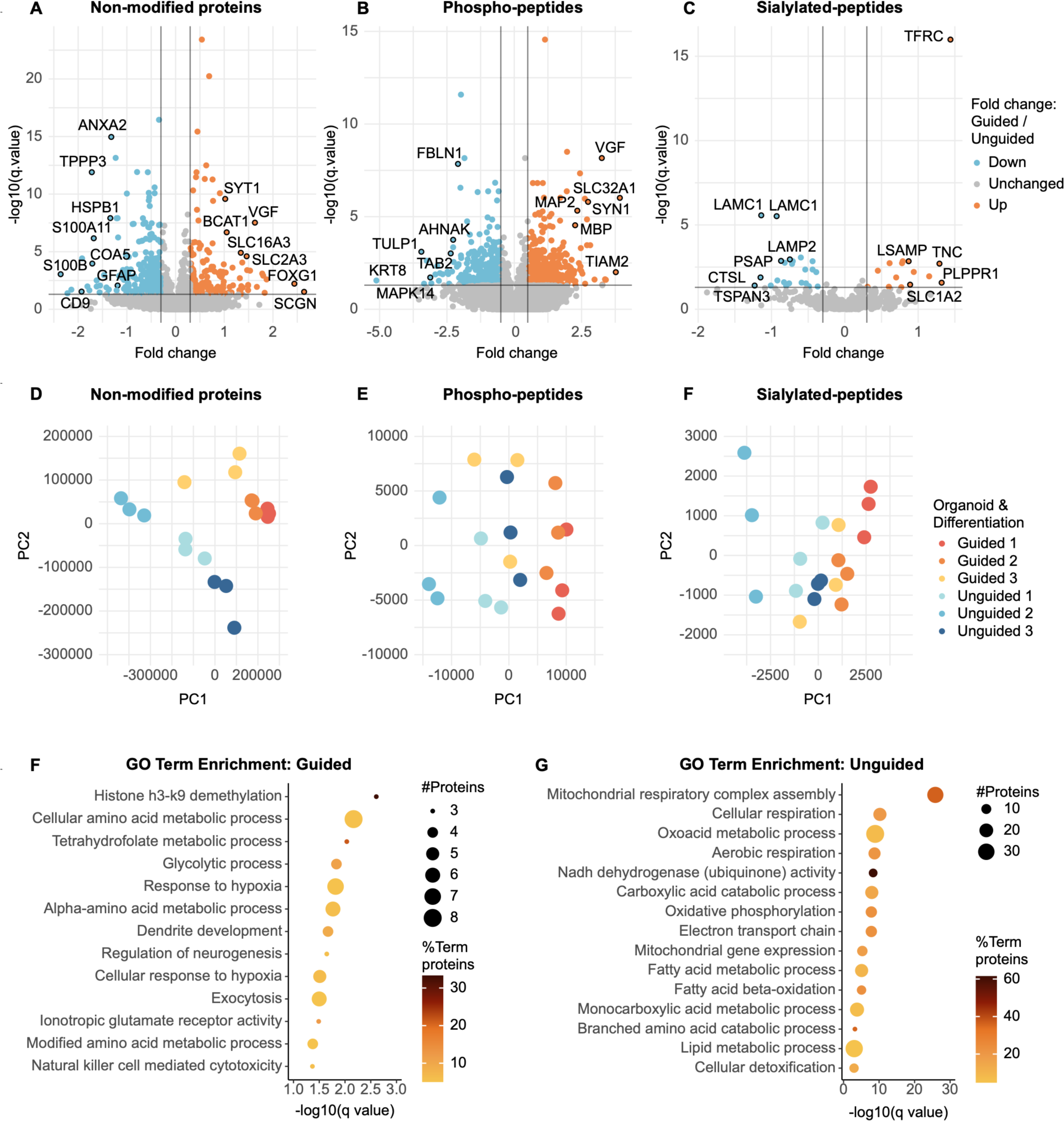
Proteomic/PTMomic profiles distinguish guided and unguided forebrain organoids (FOs) based on neuronal and metabolic proteins. (A-C) Volcano plots showing the fold change and -log10(q-value) when comparing levels of (A) non-modified proteins, (B) phospho-peptides and (C) sialylated peptides in day 40 guided vs unguided FOs using proteomics from three independent differentiations (n = 9, q≤0.05 and fold change±0.3 considered significant, Rank products test). (D-F) Principal component analysis based on (D) the non-modified proteins, (E) the phospho-peptides and (F) the sialylated peptides quantified by the proteomic analysis in guided (blue) vs unguided FOs (orange) labelled according to the originating differentiation (1-3). (G-H) GO term enrichment analysis listing the pathways that were significantly enriched (q≤0.05, fold change±0.3), amongst the non-modified proteins with (G) increased abundance and (H) decreased abundance in guided vs unguided FOs with the dot size signifying the number of significantly different proteins in the pathway and the colour indicating how many percent these constitute out of the total number of proteins in the pathway (two-sided hypergeometric test with Bonferroni step-down).

The proteins, which were significantly more abundant in guided FOs included Forkhead Box G1 (FOXG1), an essential transcription factor for brain development^33^, and other neuron-specific proteins such as synaptotagmin 1 and the neurosecretory protein VGF, whereas the glial markers glial fibrillary acidic protein (GFAP) and S100B were more abundant in unguided FOs (Fig. 2A, Table S1A-B). GO term enrichment analysis of proteins, which were significantly more abundant in guided FOs accordingly included “dendrite development”, “regulation of neurogenesis”, “exocytosis” and “ionotropic glutamate receptor activity” (Fig. 2F). The phospho-peptides that were more abundant in guided FOs also arose from numerous neuronal/synaptic proteins including VGF, Map2 and synapsin 1 (Syn1) (Fig. 2B, Table S1C-D). Proteins with significantly higher sialylation abundance in guided FOs included LSAMP, a protein that promotes neuronal growth and axon targeting, and SLC1A2, a glutamate transporter in the synaptic cleft (Fig. 2C, Table S1E-F). Resultantly, GO term enrichment analysis of this group had “axogenesis” and “neuron development” as the top terms (Fig. S2).

Surprisingly, GO term enrichment analysis of the proteins, which were significantly more abundant in unguided FOs, indicated these to be mainly related to energy metabolism with terms such as “mitochondrial respiratory complex assembly”, “Nadh dehydrogenase activity”, “oxidative phosphorylation (OXPHOS)” and “fatty acid β-oxidation” (Fig. 2G).

Correspondingly, glycolytic process was enriched amongst the significantly more abundant proteins in guided FOs (Fig. 2F). Another unexpected result was the increased abundance of sialylated lysosomal proteins including LAMP2 and prosaposin (PSAP) in unguided FOs. Overall, the proteomic analysis pointed to key differences in abundances and PTMs of neuronal and metabolic proteins between guided and unguided FOs.

### Metabolomic and lipidomic profiles differ between guided and unguided FOs

Given the difference in abundance of energy metabolism proteins, we performed LC-MS/MS-based metabolomic/lipidomic analysis of five unguided and five guided FOs differentiated in parallel. From the analysis 300 metabolites and 794 lipid-species were annotated and quantified (Table S2 and S3). The 300 identified metabolites belonged to a number of metabolic pathways with enrichment broadly of phospholipid biosynthesis, amino acid metabolism and glucose metabolism (Fig. S3). The metabolite profiles clearly separated the guided and unguided FOs on PCA (Fig. 3A-B). The lipidomic profiles also differed with significantly increased abundance of 35 lipids in guided FOs and 68 in unguided FOs (Fig. 3C). Looking more broadly at the contribution of the identified lipid classes to the total lipid amount in each FO, four of these showed significant differences with higher percentages of hexocylceramides, phosphatidylglycerols and carnitines in unguided FOs and higher percentage mainly of oleamide (in the “other” subgroup) in guided FOs (Fig. 3D, Table S3C). Oleamide is a fatty acid amide, which interacts with cannabinoid receptors and other neurotransmitter systems^34^, and can potentially stimulate neurogenesis^35^. Hexocylceramides are critical to the structure and function of myelin and enriched in oligodendrocytes^36^. Combined the metabolomic/lipidomic analysis of guided and unguided FOs supported the finding of different metabolic protein expression levels and also pointed to potential differences in the cellular composition of the FOs.

**Figure 3:**
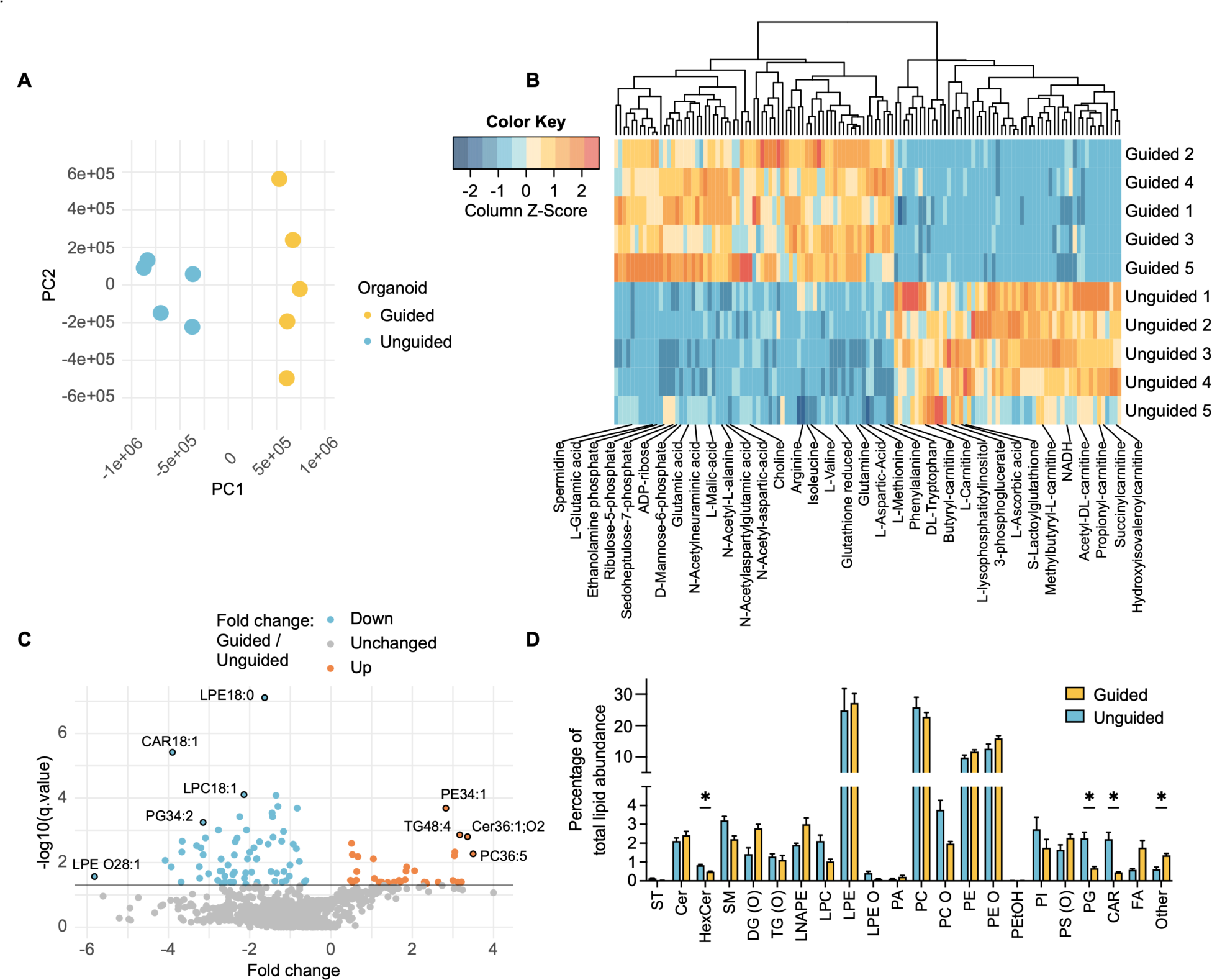
Guided and unguided forebrain organoids (FOs) have different metabolomic/lipidomic profiles. (A) Principal component analysis based on the metabolomic data in guided (blue) vs unguided (orange) FOs (n=5). (B) Heatmap of metabolites with significantly different (q≤0.05) abundances in guided vs unguided FOs ordered by hierarchical clustering (n=5, Rank products test). (C) Volcano plots of the lipidomic data showing the fold change and -log10(q-value) when comparing guided vs unguided FOs (n = 5, q≤0.05 considered significant, Rank products test). (D) Identified and annotated lipids sorted in lipid classes with levels of each lipid class shown as percentage of total lipid abundance in each organoid. Mean ± SEM. *q≤0.05 (n=5, Student’s t-test with Benjamini-Hochberg correction for multiple testing).

### Increased neuronal content in guided FOs compared to unguided

To explore the enrichment of neuron-related terms in proteins of increased abundance in guided FOs, we selected all proteins related to synapses and/or neurodevelopment amongst them (Fig. 4A). The network created from these contained glutamate receptors (GRIK3, GRIA2), important proteins in GABA signalling (GAD1, GAD2, SLC32A), brain-derived neurotrophic factor (BDNF) and key transcription factors in brain development (FOXG1, PAX6 and POU3F3 (BRN1)). GO term enrichment of the proteins with increased phosphorylation levels in guided FOs further supported that generation of neurons and neuronal projection was enhanced in guided FOs (Fig. 4B). To confirm this, we performed western blotting for Syn1 and Map2, finding significantly increased levels of both in day 40 guided FOs (Fig. 4C-D). Interestingly, these differences persisted in day 120 FOs (Fig. 4E-F). Accordingly, levels of four out of five neurotransmitters identified by the metabolomic analysis, including glutamate (glutamic acid), were significantly increased in guided FOs (Fig. 4G). To evaluate if these differences resulted in different numbers of functional synapses, we performed ICC and examined co-localisation of the presynaptic protein Syn1 and postsynaptic density protein 95 (Psd95). At day 40 the guided FOs appeared to have higher expression and colocalization of Syn1/Psd95, however at day 86 the levels were comparable (Fig. 4H). Multi electrode array (MEA) recordings of the spontaneous electrophysiological activity of day 120 guided and unguided FOs did not reveal any significant differences in frequency or amplitude of spikes (Fig. 4I-J). We thus found substantial evidence for increased neuronal content in guided FOs both at early and later time points, likely because the guided differentiation approach “forces” the FOs to generate mature neurons in larger quantities earlier during the differentiation.

**Figure 4:**
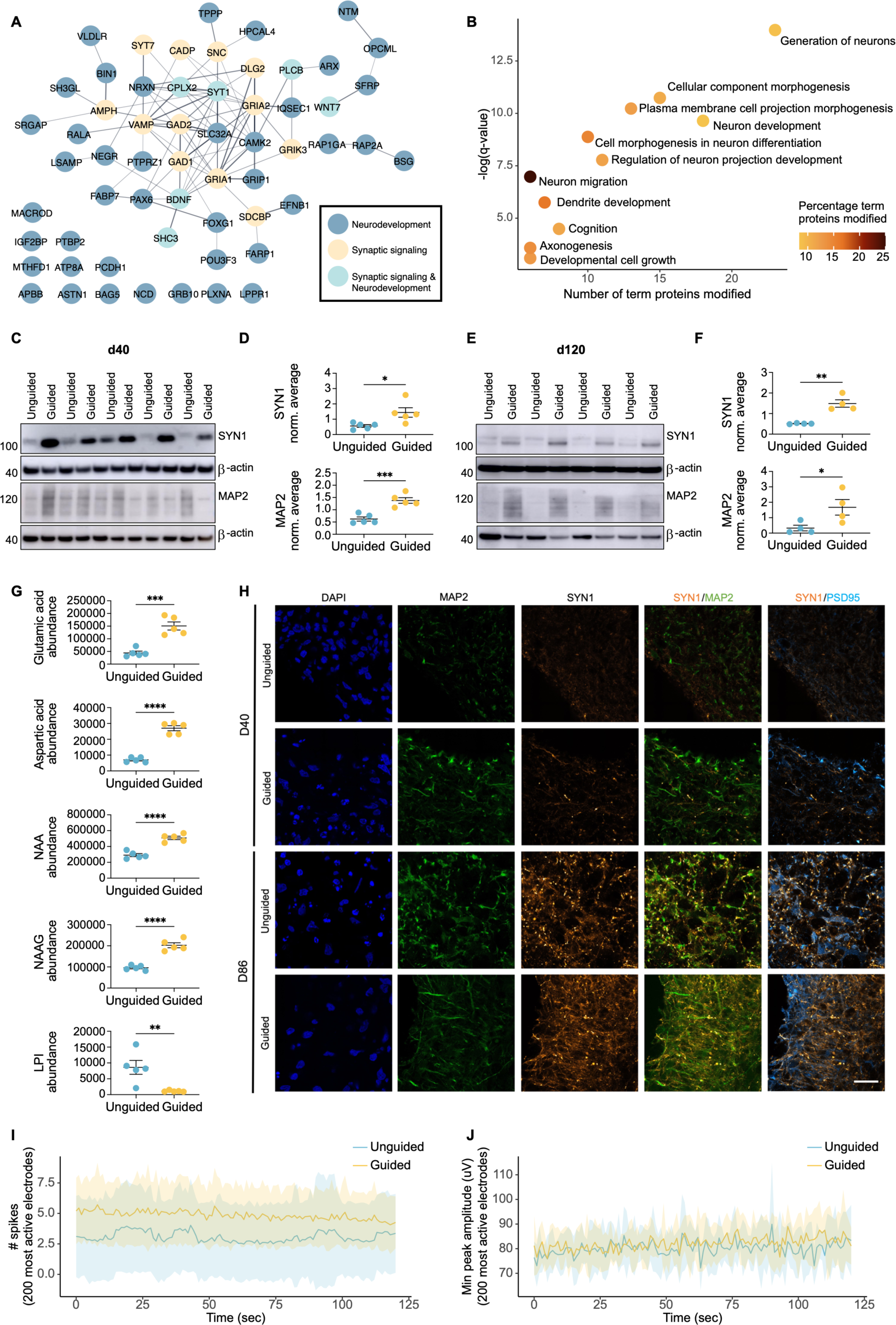
Higher abundance of neuronal and synaptic proteins in early and late stage guided forebrain organoids (FOs) (A) String network of proteins involved in synaptic signalling and/or neurodevelopment of significantly increased abundance (q≤0.05, fold change±0.3) in guided vs unguided FOs in the proteomic analysis (n=9, Rank products test). (B) GO term enrichment analysis of the pathways that were significantly enriched, amongst the proteins with significantly increased (q-value≤0.05, fold change±0.3) abundance of phospho-peptides in guided vs unguided FOs based on the number of significantly different proteins in the pathway. Dot colour indicates how many percent these constitute out of the total number of proteins in the pathway (two-sided hypergeometric test with Bonferroni step-down). (C-F) Representative western blots and quantification of synapsin 1 (SYN1) and microtubule associated protein 2 (MAP2) levels in (C-D) day 40 and (E-F) day 120 guided and unguided FOs. Protein expression levels normalised to β-actin and the average of all samples in each blot. Mean ± SEM, *p≤0.05, **p≤0.01, ***p≤0.001 (Student’s T-test). (G) Abundance levels of neurotransmitters quantified by metabolomics. NAA = N-acetylaspartate, NAAG = N-acetylaspartylglutamate, LPI = Lysophosphatidylinositol. Mean ± SEM, **p≤0.01, ***p≤0.001, ****p≤0.0001 (n=5, Rank products test). (H) Immunocytochemistry of day 40 and 86 guided and unguided FOs for DAPI (dark blue), microtubule associated protein 2 (Map2, orange), Synapsin 1 (Syn1, green) and postsynaptic density protein 95 (Psd95, light blue). Scalebar = 20 μm. (I-J) Multi electrode array recording from the 200 most active electrodes from day 120 guided and unguided FOs from two independent differentiations showing the number of spikes and minimum peak amplitude in μV over 2 min. Mean ± SEM (n=5-6).

### Increased radial glia/astrocytic content with cytoplasmic FOXG1 localization in unguided FOs

As the proteomic analysis had identified increased levels of GFAP and S100B in unguided FOs, we hypothesised that the increased neuronal content in guided FOs were at the expense of decreased radial glia and/or astrocyte content. GFAP and S100B are markers of radial glia and astrocytes, which are likely present at the later time points examined here.

ICC for GFAP at day 40 and 86 showed lower expression levels at both time points (Fig. 5A-B), which was confirmed by western blotting for GFAP at day 40 and 120 where a significant substantial difference was seen (Fig. 5C-D). Surprisingly, the subcellular localisation of FOXG1 in GFAP^+^ cells differed markedly between the two FO types at day 86. In unguided FOs FOXG1 co-localised with the cytoplasmic GFAP staining, whilst in guided FOs FOXG1 was nuclear (Fig. 5B). The cytoplasmic FOXG1 expression was not seen in the neuronal population as identified by Map2 (Fig. 5B), indicating that this transition was specific for the GFAP^+^ cells. As FOXG1 is nuclear in progenitor cells, but cytoplasmic in differentiating cells^37^, this might indicate a difference in the differentiation stage of GFAP^+^ cells between guided and unguided FOs.

**Figure 5:**
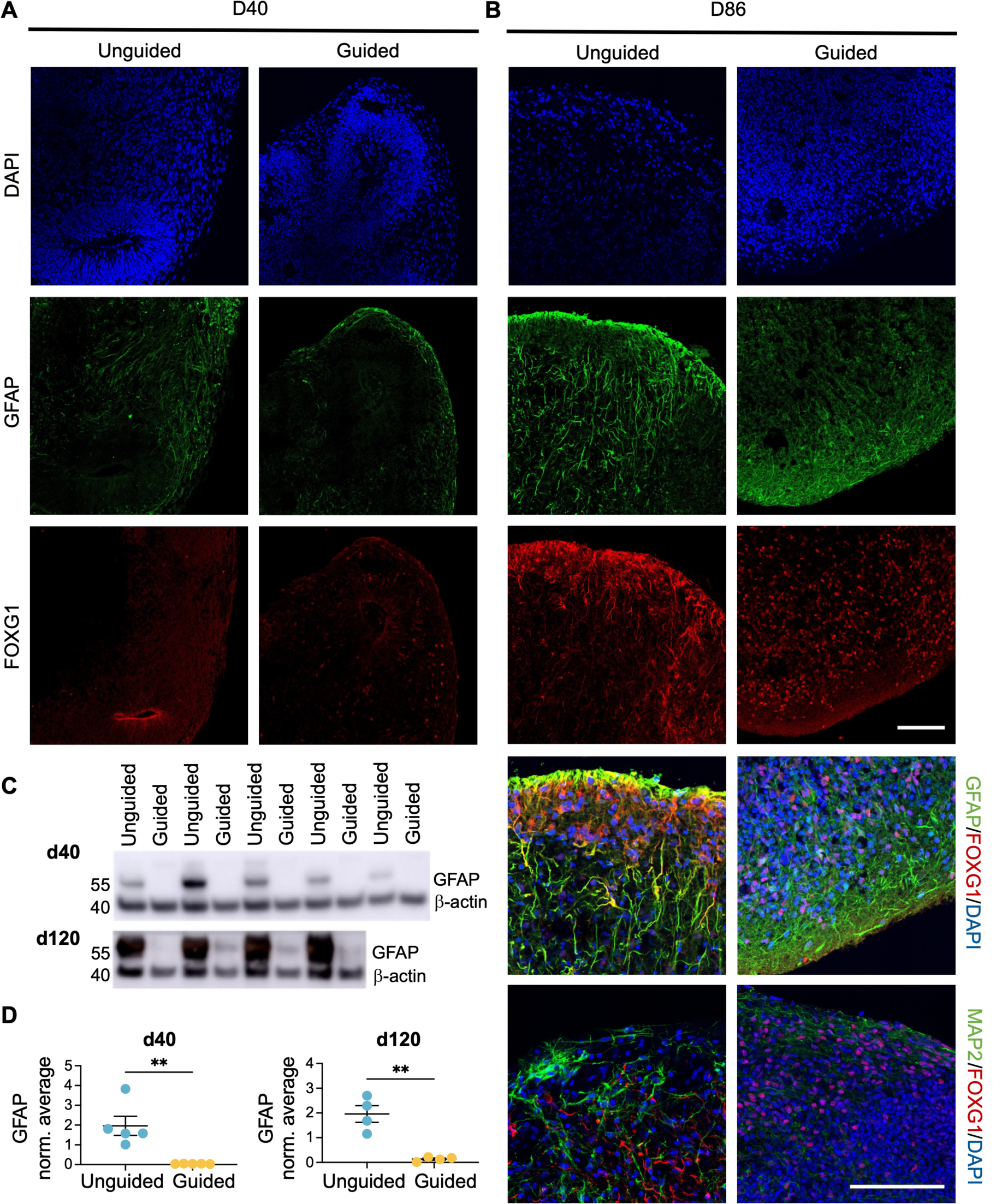
Higher abundance of GFAP+ cells with altered subcellular FOXG1 localisation in unguided forebrain organoids (FOs) (A-B) Immunocytochemistry of guided and unguided FOs at (A) day 40 and (B) day 86 for DAPI (dark blue), Forkhead Box G1 (Foxg1, red), glial fibrillary acidic protein (Gfap, green) or microtubule associated protein 2 (Map2, green). Scalebars = 100 μm. (C-D) Representative western blots and quantification of GFAP levels in day 40 and day 120 guided and unguided FOs. Protein expression levels normalised to β-actin and the average of all samples in each blot. Mean ± SEM, **p≤0.01 (n=4/5, Student’s T-test).

Overall, this indicated a substantial difference in the amount of GFAP^+^ cells between guided and unguided FOs.

### Increased mitochondrial content and OXPHOS proteins in unguided FOs

Perhaps related to these differences in the cell composition of the FOs, the proteomic analysis had identified marked differences in abundances of energy metabolism-related proteins between guided and unguided FOs. Numerous proteins of the TCA cycle, complex I, fatty acid β-oxidation and mitochondrial proteins were found in significantly increased abundance in unguided FOs, whilst in guided FOs glycolysis proteins were significantly increased (Fig. 6A). Interestingly, the metabolomic analysis identified significantly increased levels of glycolysis/penthose phosphate pathway metabolites in guided FOs (Fig. 6B). Corresponding with the increase in fatty acid β-oxidation proteins, unguided FOs had significantly elevated levels of various carnitines (Fig. 3B, 6B). The main function of carnitines is to transfer long-chain fatty acids to mitochondria for β-oxidation^38^.

**Figure 6:**
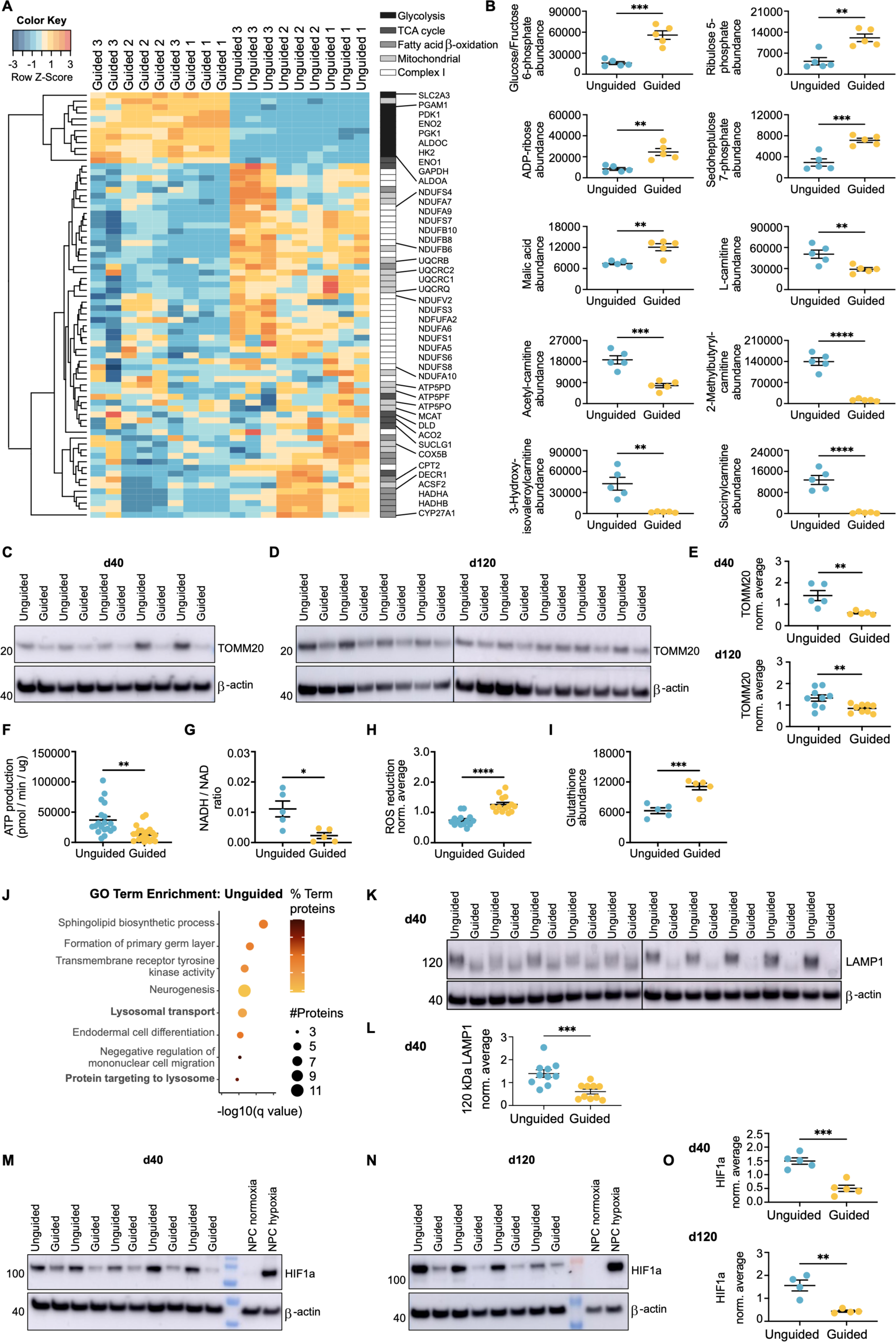
Increased levels of mitochondria and OXPHOS in unguided vs unguided forebrain organoids (FOs) (A) Heatmap of non-modified proteins related to glycolysis, TCA cycle, fatty acid β-oxidation, mitochondria and complex I with significantly different (q-value≤0.05, fold change±0.3) abundances in guided vs unguided FOs ordered by hierarchical clustering (n = 9, Rank products test). (B) Abundance levels of metabolites related to glycolysis/pentose phosphate pathway or fatty acid β-oxidation quantified by metabolomics. Mean ± SEM, **p≤0.01, ***p≤0.001, ****p≤0.0001 (Rank products test). (C-E) Representative western blots and quantification of TOMM20 levels in (C,E) day 40 and (D,E) day 120 guided and unguided FOs. Protein expression levels normalised to β-actin and the average of all samples in each blot. Mean ± SEM, **p≤0.01 (n=5/9, Student’s T-test). (F) ATP production in pmol/min normalised to protein content (μg) measured by Seahorse on sectioned day 80 guided and unguided FOs from 3 independent differentiations. Mean ± SEM, **p≤0.01 (n=20, Student’s T-test). (G) Ratio of NADH/NAD as quantified by metabolomics. Mean ± SEM, *p≤0.05 (n=5, Rank products test). (H) Reduction of reactive oxygen species (ROS) by guided and unguided FOs from three independent differentiations, relative fluorescent units normalised to average of all samples per differentiation. Mean ± SEM, ****p≤0.0001 (n=14, Student’s T-test). (I) Abundance levels of reduced glutathione as quantified by metabolomics. Mean ± SEM, ***p≤0.001 (n=3, Rank products test). (J) GO term enrichment analysis listing the pathways that were significantly enriched (q≤0.05), amongst the proteins with increased sialylation levels in unguided vs guided FOs with the dot size signifying the number of significantly different proteins in the pathway and the colour indicating how many percent these constitute out of the total number of proteins in the pathway (two-sided hypergeometric test with Bonferroni step-down). (K-L) Representative western blots and quantification of glycosylated 120 kDa LAMP1 levels in day 40 guided and unguided FOs. Protein expression levels normalised to β-actin and the average of all samples in each blot. Mean ± SEM, ***p≤0.001 (n=10, Student’s T-test). (M-O) Representative western blots and quantification of HIF1α levels in (M,O) day 40 and (N,O) day 120 guided and unguided FOs. Neural precursor cells (NPCs) cultured in 2D under normoxic (20% oxygen) or hypoxic (1% oxygen) conditions for 4 hours as controls. Protein expression levels normalised to β-actin and the average of all samples in each blot. Mean ± SEM, **p≤0.01, ***p≤0.001 (n=4/5, Student’s T-test).

To address whether the increase in OXPHOS-related proteins was caused by enhanced mitochondrial content in the unguided FOs, we examined levels of the mitochondrial marker TOMM20. ICC at day 40 indicated a small, but significant increase in TOMM20 expression in unguided FOs (Fig. S4) and western blotting showed significantly elevated levels of TOMM20 at both day 40 and 120 (Fig. 6C-E), supporting that unguided FOs contain relatively more mitochondria. In line with this, the total ATP production, as measured by Seahorse analysis, was significantly higher in unguided FOs (Fig. 6F). This correlated with a significantly larger NADH/NAD ratio in the unguided FOs as identified by the metabolomic analysis (Fig. 6G). Perhaps due to higher OXPHOS levels in unguided FOs, the ability to reduce reactive oxygen species (ROS) and the levels of reduced glutathione were significantly decreased compared to guided FOs (Fig. 6H-I).

Combined, our results indicated that the unguided FOs had increased mitochondrial content and relied more heavily on OXPHOS than the guided FOs, which were utilising glycolysis more.

### Glycolysis protein levels do not correlate with HIF1α levels in guided and unguided Fos

The finding of increased mitochondrial content and OXPHOS in unguided FOs seemed counterintuitive in light of the results indicating accelerated neuronal differentiation in guided FOs. During normal differentiation from neural precursors to postmitotic neurons, a metabolic switch from glycolysis to OXPHOS happens^39,40^. If more neurons of increased maturity were present in guided than unguided FOs, this should result in relatively more mitochondria and OXPHOS in guided FOs. Besides the difference in mitochondrial content, we also observed changes implying different lysosomal content. Proteins involved in “lysosomal transport” and “protein targeting to lysosomes” had significantly increased sialylation levels in unguided FOs (Fig. 6J). Western blotting for the glycosylated form of LAMP1, a lysosomal marker, showed significantly increased abundance at day 40, confirming the proteomics data (Fig. 6K-L). Lysosomal proteins such as LAMP1 require n-linked glycosylation, including sialylation, for proper targeting to and function in lysosomes^41^ and their upregulation likely indicate increased lysosomal content in unguided FOs.

The guided FOs had significantly higher levels of monocarboxylate transporter 4 (MCT4) and hexokinase 2 (HK2) (Table S1), which are upregulated in highly glycolytic cells^42^. Based on the increased abundance of these and other glycolytic proteins, the GO term enrichment of guided FO proteins included “response to hypoxia” (Fig. 2F). As hypoxia-inducible factor 1α (HIF1α) can upregulate expression of HK2, MCT4 and other glycolytic proteins^42^, we aimed to test whether differences in hypoxia levels in the FOs could be causing the contrasting metabolic profiles. However, consistent with the significantly larger size, HIF1α levels were significantly enhanced in unguided FOs at day 40 and 120 (Fig. M-O). Differences in hypoxia could therefore not explain the discrepancy in OXPHOS/glycolysis-reliance between guided and unguided FOs.

### Differences in cellular composition of guided and unguided FOs

To understand whether the observed dissimilarities in neuronal and metabolic content arose from differences in the cellular composition of the FOs, we performed single cell RNA sequencing (scRNAseq) on a total of 9913 cells from day 20 and day 40 guided and unguided FOs (n = 3 per group) (Fig. S5A-C). Clustering and differential gene expression analysis resulted in 12 clusters of progenitors and neurons, resembling the main populations found in early forebrain development and comparable to earlier scRNAseq analyses of FOs (Fig.7A-B, Fig. S6, Table S4)^16,17,28^. Separating the cells on time points and organoid type (Fig. 7C-E), revealed that markers of cortical hem (CH) (Fig. 7D; LMX1A) and choroid plexus (ChP) (Fig. 7D; TTR) and the corresponding cell clusters were found at noticeably higher levels in unguided FOs at day 40 (Fig. 7C-E). Correspondingly, reelin, a key marker of Cajal Retzius cells (CR), which mainly arise from cortical hem^31^, were more highly expressed in unguided FOs (Fig. D; RELN). Surprisingly, the cell populations expressing markers of medial ganglionic eminence (mGE) and interneurons (INs) (Table S4) were found only in guided FOs as demonstrated by markers for migrating (Fig. 7D; DLX2) and mature interneurons (Fig. 7D; GAD2), which were almost exclusively expressed in guided FOs.

**Figure 7:**
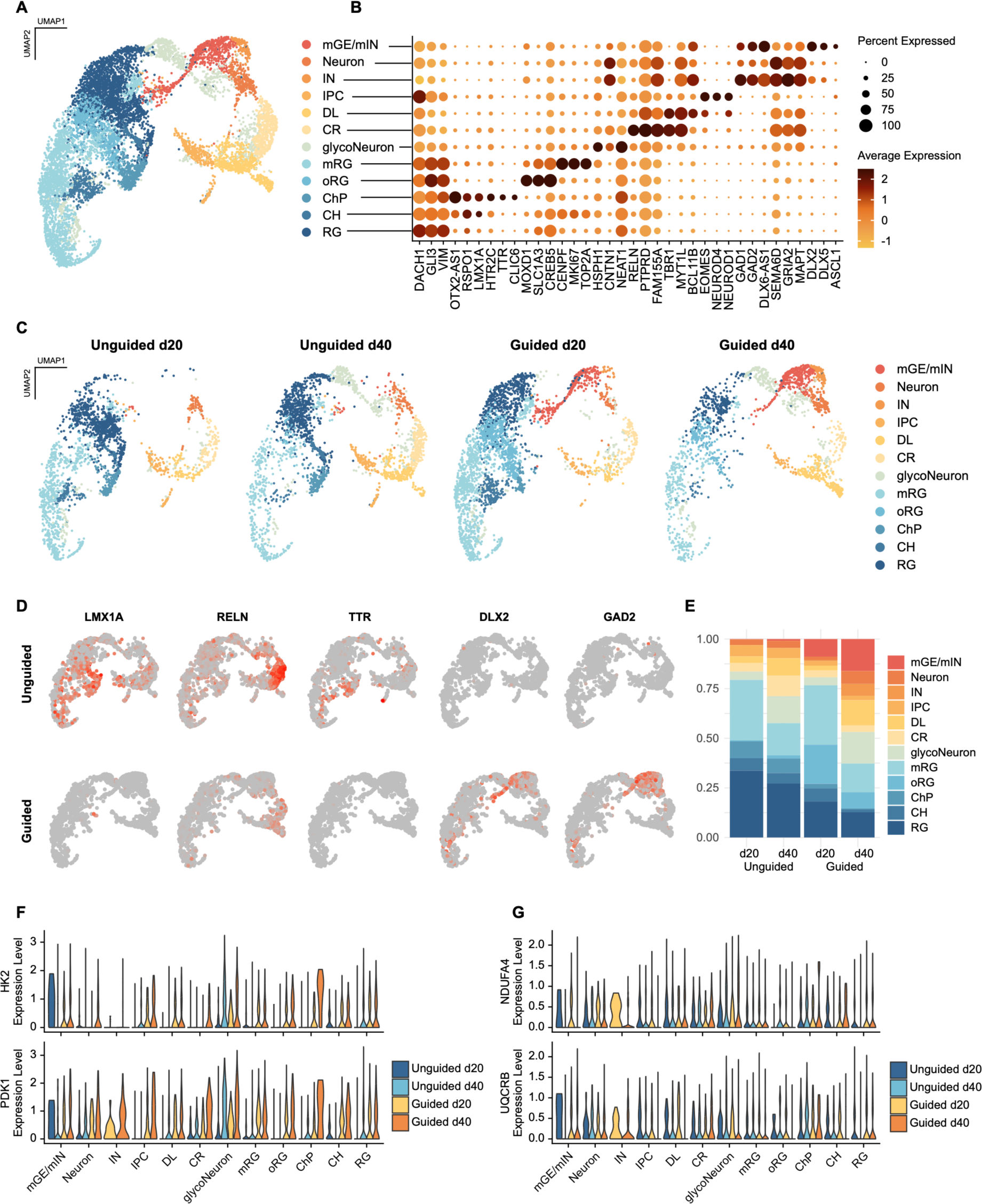
scRNAseq reveal differences in cellular composition and expression of glycolytic markers in unguided vs unguided forebrain organoids (FOs) (A) UMAP of single cell RNA sequencing (scRNAseq) data from day 20 and 40 guided and unguided FOs (n=3 per time point), analysed by splitting the dataset into the four conditions, normalising and identifying variable features of each dataset before reintegrating based on repeatedly variable features. This identified the following 12 clusters: CR; Cajal Retzius cells, RG; radial glia, oRG; outer RG, mRG; mitotic RG; CH; cortical hem, ChP; choroid plexus, DL; deep layer neurons, IPC; intermediate progenitors, IN; interneurons, mGE/mIN; medial ganglionic eminence/migratory INs. (B) Expression of key markers for each of the 12 clusters with colour indicating average expressen levels and dot size depicting percentage of cells expressing the marker. (C) UMAP split based on cell origin; unguided or guided, day 20 and day 40 with similar cluster identities as above. (D) Feature plots showing expression levels for markers of cortical hem (LMX1A), Cajal Retzius cells (RELN), choroid plexus (TTR) and interneurons (DLX2, GAD2) in day 40 unguided and guided FOs. (E) Proportions of each cell type in day 20 and 40 guided and unguided FOs. (F-G) Violin plots of RNA expression levels for key glycolysis enzymes, Hexokinase 2 (HK2) and Pyruvate Dehydrogenase Kinase 1 (PDK1), and OXPHOS proteins, NDUFA4 and UQCRB, across the different cell types.

However, the proteomic analysis had identified GAD1 and GAD2 protein expression in unguided FOs albeit at significantly lower levels than in guided FOs (Table S1B). In accordance with the proteomic findings, GFAP was more widely expressed in unguided FOs and FOXG1 levels were higher in guided FOs (Fig. S5D). Given that FOXG1 is more highly expressed in the ventral forebrain than the dorsal, this is consistent with the increased numbers of inhibitory neurons in the guided organoids^43^.

Both organoid types at day 40 contained a population of neurons with high expression of glycolytic genes and stress markers such as HSPH1 and NEAT1 (Fig. 7B,F, Table S4). To determine differences in the developmental trajectories we performed pseudotime analysis on the scRNAseq data from both organoid types separately (Fig. S7A-D). This supported a unique lineage from mitotic radial glia via mGE to migratory and mature interneurons in guided FOs (Fig. S7A-B). In unguided FOs a trajectory from radial glia (RG) through intermediate progenitors (IPCs) to Cajal Retzius and deep layer neurons, corresponding to indirect neurogenesis, was identified (Fig. S7C-D). This was not seen in guided FOs where the RG population connecting to IPCs was not present (Fig. S5A-B). This might indicate that formation of IPCs happens prior to day 20 in guided FOs as the linage from IPCs to Cajal Retzius and deep layer neurons remained. In both organoid types the glycolytic neurons (glycoNeuron) appeared to be generated through direct neurogenesis from RG (Fig. S7A-D). The proportion of glycoNeurons in guided and unguided FOs was not markedly different and as such could not explain the higher levels of glycolytic proteins in guided FOs. Expression of key glycolysis proteins HK2 and PDK1 appeared universally increased in guided FOs at day 40 across cell types (Fig. 7F). In contrast, levels of OXPHOS-related transcripts, NDUFA4 and UQCRB, were more uniform between the four conditions and mainly varied between cell types with the highest expression in ChP and glycoNeurons (Fig. 7G). Given the increased expression of OXPHOS transcripts in ChP, the presence of this cell population in mainly unguided FOs could contribute to their increased OXPHOS. In support of this, transcripts for PGC-1α (PPARGC1A), a master regulator of mitochondrial biogenesis, were at the highest levels in ChP in day 40 unguided FOs (Fig. S5E)^39^.

Overall, the scRNAseq identified differences in the cellular content, which could partly explain the observed metabolic differences, but also supported a general increase in glycolysis transcripts across cell types in guided FOs.

## Discussion

In the present study we aimed to identify how guided and unguided FOs differ to enable more informed decisions on their applications. To achieve this we generated FOs by both approaches in parallel starting from the same batches of iPSCs and applied an unbiased multi-omics approach to quantify the resulting differences.

Morphologically the unguided FOs showed more variability (Fig. S1) due to the larger neuro-ventricular structures, which varied in shape and size. Examining the overall protein composition (Fig. 2D) the inter-organoid variability was correspondingly larger between unguided than guided FOs, although not substantially. In both cases, batch-to-batch variability between differentiations was observed and appeared the largest source of variation. The unguided FOs were significantly larger at day 40 despite arising from a smaller number of iPSC. This could be caused by the addition of extrinsic signalling molecules, which stimulate a more rapid neuronal specification and differentiation and less proliferation in guided FOs. However, the use of agitation culture for unguided FOs starting from day 10 might also lead to enhanced growth through increased flow of nutrients.

The promotion of neuronal differentiation in guided FOs led to a long-term increase in neuronal and synaptic proteins and neurotransmitter levels. This did not translate into increased synapse formation or activity at the later time points, although we cannot exclude that more high-resolution techniques than applied here might be needed to detect potential differences. The increased neuronal content in guided FOs was countered by a substantial increase in glial progenitor/astocytic content in unguided FOs as judged by GFAP/S100B levels. This difference in the proportion of neuronal to glial content could be caused by differences in FOXG1 levels, which were significantly higher in guided FOs. FOXG1 is a critical factor for directing neuronal identity and overexpression of FOXG1 in neural precursors *in vitro* increases the ratio of neural to glial progenitors^33^. Interestingly, in unguided FOs, a cytoplasmic translocation of FOXG1 was observed at day 86 in areas with GFAP+ cells, specifically. As FOXG1 is nuclear in progenitor cells but as a result of FGF2 signalling cytoplasmic in differentiating cells^37^, this could indicate increased differentiation of radial glia progenitors to astrocytes in unguided FOs. The choroid plexus is known to produce and secrete FGF2, which can shift the progeny fate of multipotent cortical progenitors towards glial lineage^44,45^. This could suggest that intra-organoid signalling from choroid plexus in unguided FOs could affect the lineage determination of cortical progenitors leading to increased differentiation of glial cells.

Surprisingly, the scRNAseq analysis revealed a marked difference in the neuronal cell type composition as the unguided FOs contained almost no interneurons. This is in contrast to earlier studies, which have identified interneurons in unguided FOs^1,16^. However, the FOs applied in the current study were generated with commercial kits with media compositions that, although based on the published methods^5,17^, are undisclosed. If they differ from the published methods this might explain this discrepancy. As the proteomic analysis, which was done on differentiations independent from the scRNAseq, had identified GAD1 and GAD2 expression in both organoid types (1.4- and 0.8-fold higher in guided FOs, respectively) this indicates that the interneuron contribution to unguided FOs can vary significantly between differentiations with this method.

The findings pointing to increased OXPHOS and mitochondrial content in unguided FOs were also surprising at first as this could be consistent with enhanced neuronal differentiation. A metabolic shift from glycolysis dependence to OXPHOS and increasing mitochondrial mass is required for neuronal differentiation^46,47^. This increase in mitochondrial mass, which is necessary for neuronal commitment, is orchestrated by PI3K/mTOR signalling, PGC-1α and TFAM, which are master regulators of mitochondrial biogenesis^39,48^. Given that they can also promote lysosomal biogenesis this could explain the concomitant increased mitochondrial and lysosomal abundance in unguided FOs.

However, an increase in glycolysis and pentose phosphate pathway (PPP) metabolites is also seen during neuronal differentiation^39^. Aerobic glycolysis and PPP is hypothesised to support the biosynthesis required for synapse and neurite formation^39,49^. As such the enhanced levels of glycolytic/PPP proteins and metabolites are consistent with the higher protein expression of neuronal/synaptic proteins in guided FOs as well as higher levels of reduced glutathione, which the PPP is key in maintaining by providing NADPH^39^. However, this does not explain the increased mitochondrial content in unguided FOs or the enrichment of fatty acid beta-oxidation enzymes and metabolites.

One explanation for this could be the higher levels of GFAP+ radial glia/astrocytes in unguided FOs. Although the prevailing hypothesis is that neurons are mostly oxidative and astrocytes glycolytic, evidence of fatty acid beta-oxidation in astrocytes have recently emerged^50,51^. Astrocyte mitochondria are enriched with enzymes involved in mitochondrial β-oxidation of fatty acids and metabolise long chain fatty acids more efficiently than neuronal mitochondria^50,51^. However, studies also suggest that fatty acid β-oxidation is key for neural stem cell self-renewal^52,53^. As such the larger proportion of radial glia in day 40 unguided FOs determined by scRNAseq, and supported by increased GFAP levels, could also be a potential source of the increased fatty acid β-oxidation enzymes and metabolites. Similarly, the increased amount of choroid plexus cells in unguided FOs likely contribute to higher mitochondrial content as indicated by higher expression of PGC-1α transcripts. ChP cells are known to contain a large proportion of mitochondria due to high energy demand related to their secretory function^54^.

Conversely, the scRNAseq demonstrated increased expression of glycolytic transcripts universally across cell types in guided FOs at day 40. We therefore explored whether more pronounced hypoxia in guided FOs could be responsible for the increased abundance of glycolytic proteins as several of these are upregulated in response to hypoxia. These include MCT4, HK2 and PDK1, which inhibits pyruvate dehydrogenase activity, and thereby regulates metabolite flux through the tricarboxylic acid cycle, down-regulating aerobic respiration^42,55,56^. Yet, at day 40 and 120 HIF1α levels were significantly higher in unguided FO. This could simply be a result of their relatively larger size at these time points or it could stem from the higher abundance of mitochondria and OXPHOS, which through increased oxygen demand result in intracellular hypoxia and HIF1α stabilisation^57^. Interestingly, hypoxia can promote astrocytic differentiation of neural precursor cells through epigenetic regulation of GFAP^58^, which could be yet another factor contributing to the increased GFAP levels in unguided FOs. The effect of agitation culture on the metabolism of FOs might also contribute to differences.

In both organoid types, we identified a population of neurons with high expression of hypoxia-, glycolysis- and cellular stress markers (glycoNeurons). These neurons appeared to arise directly from radial glia, a process resembling direct neurogenesis. Whether these directly generated neurons are more susceptible to hypoxia and cell stress or they are a result of premature differentiation of neural progenitors caused by hypoxia, remains to be explored^28,59^.

In conclusion, we conducted a thorough multi-omics analysis-based comparison of early differentiation stages in guided and unguided FOs, applying two protocols which are widely used in the field. The results provide a significant resource for the field and can aid in determining the most appropriate model for certain applications. Our findings highlight significant differences in neuronal differentiation, cellular content and metabolic activity and furthermore underline the need for additional characterization of organoid models in relation to *in vivo* human brain development, to improve the accuracy of the models for *in vitro* studies of brain development and disease.

## Methods

### iPSC maintenance and forebrain organoid (FO) differentiation

The human iPSC line IMR90 clone 4 was purchased from WiCell and maintained on growth factor reduced Matrigel-coated plates (Corning) in mTESR1 medium (Stem Cell Technologies). Passaging was performed using Gentle cell dissociation reagent (Stem Cell Technologies).

Unguided FOs were generated with the Cerebral Organoid Kit (Stem Cell Technologies) according to manufacturer instructions with the following modifications: On day 0, 600,000 iPSCs per well were seeded in Aggrewell^TM^800 24-well plates (Stem Cell Technologies) pre-treated with Anti-Adherence Rinsing Solution (Stem Cell Technologies) in embryoid body (EB) Seeding medium with 50 μM ROCK inhibitor (Y27632, Stem Cell Technologies). On day 7 EBs were transferred from Aggrewell^TM^800 24-well plates using wide-bore p1000 pipette tips and embedded in Matrigel by gently mixing around 30 EBs in 100 μl Expansion medium with 150 μl cold Matrigel (Corning) using wide-bore p200 tips and plating the mixture in a circle (diameter around 10 mm) in 60 mm culture dishes. Following 30 min incubation at 37°C to polymerize the Matrigel, 4 ml Expansion medium was added per dish. On day 10 the dishes were moved to an INFORS HT Celltron orbital shaker set to 57 rpm in a 37°C incubator. On day 15, the FOs were released from the Matrigel by gentle pipetting with wide-bore p1000 tips and around 16 FOs per dish cultured in 5 ml medium.

Guided FOs were generated using the STEMdiff Dorsal FO Differentiation Kit (Stem Cell Technologies) and maintained with the STEMdiff Neural Organoid Maintenance Kit (Stem Cell Technologies) according to manufacturer protocols with the following modifications: during the expansion period guided FOs were cultured in low-binding 24-well plates with 4-5 FOs per well to minimise fusion of the FOs. At around day 20, the FOs were moved to 60 mm low-binding culture dishes with 16 FOs per dish.

For both FO differentiations 1% Penicillin/Streptomycin (Pen/Strep, Gibco) and 1 μg/ml Amphotericin B (Amp B, Thermo Scientific) was added to the medium from day 10 onwards. The number of FOs per dish was gradually decreased from 16 to 5 with increasing size of the FOs.

### Freezing, cryosectioning, immunocytochemistry and imaging

FOs from 2 or 3 independent differentiations were fixed using 4% paraformaldehyde (Thermo Scientific) in PBS (Gibco) for 1 hour at room temperature (RT) and subsequently soaked in 30% sucrose at 4 °C for at least one night in PBS and until embedding. The FOs were embedded in OCT mounting medium, frozen in ethanol at below -50 °C and stored at - 70°C. FOs were sectioned on a Leica CM1860 cryostat at -17 to -20 °C in 30 µm sections and plated on glass slides. The slides were stored at -20 °C.

The slices were hydrated with PBS, washed 2x 5 min in 0.1% TritonX100 (Plusone) in PBS and incubated for 30 min with permeabilization and blocking buffer (5% donkey or goat serum and 0.1% TritonX100 in PBS). The slices were incubated overnight (ON) at 4 °C with primary antibodies in 1% serum and 0.1% TritonX100 in PBS. The samples were washed 3x 10 min with 0.1% TritonX100 in PBS and incubated with secondary antibodies for 1h in the dark at RT. The slices were washed 3x 10 min with PBS and mounted with DAPI-containing ProLong Diamond Antifade mountant (Invitrogen). Slices were kept at 4 °C in the dark until image acquisition.

Images were aquired using a Nikon A1R confocal unit on a Ti-2 LFOV microscope with equal settings between the two organoid types. The images shown were edited using ImageJ version 1.53t. Quantification of TOMM20 ICC labelling was performed using the analyse particles function in ImageJ^60^.

| **Antigen** | **Species** | **Company** | **Cat. No.** | **Dilution** |
| --- | --- | --- | --- | --- |
| Ctip2 | rat | Abcam | ab18465 | 1:500 |
| Dcx | rabbit | Cell Signaling | 4604 | 1:800 |
| Foxg1 | rabbit | Thermo | 702554 | 1:250 |
| Gfap | mouse | Invitrogen | MA515086 | 1:500 |
| Map2 | mouse | Sigma | M1406 | 1:2000 |
| Ncad | mouse | Cell Signaling | 14215 | 1:500 |
| NeuN | rabbit | Cell Signaling | 24307 | 1:50 |
| Psd95 | goat | Abcam | ab12093 | 1:100 |
| Satb2 | rabbit | Abcam | ab34735 | 1:500 |
| Syn1 | rabbit | Abcam | ab254349 | 1:500 |
| Tbr1 | rabbit | Abcam | ab183032 | 1:100 |
| Tomm20 | rabbit | Abcam | ab186735 | 1:250 |

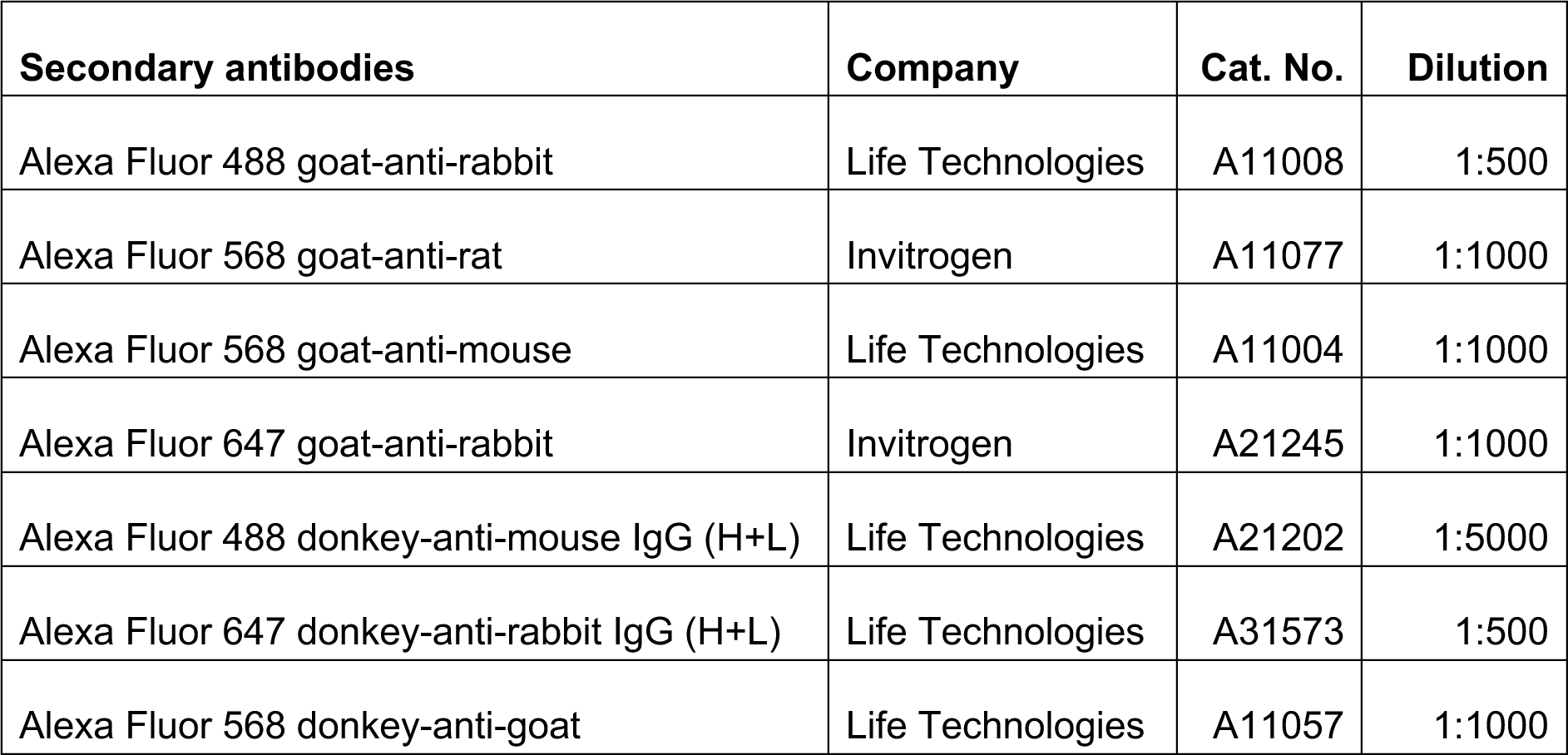

### Protein lysis, digestion and TMT-labelling for proteomic analysis

Day 40 FOs were transferred to low-bind eppendorf tubes, washed x1 with PBS and snap-frozen on dry ice. FOs were lysed in 1% SDC (Sigma) in 50 mM TEAB (Sigma) with 10 mM DTT (Sigma), pH8, and sonicated 2 x 10 sec on ice at 35% amplitude. Following sonication, samples were centrifuged at 5000 rcf for 10 min at RT and supernatant transferred to new Eppendorf tubes. Protein concentration was determined by Nanodrop (Implen). 40 μg protein per sample was alkylated for 30 min with 20 mM Iodoacetamide (Sigma) and digested with 5% trypsin (w/w) for 4 hours at 37°C. The samples were alkalised with 1 M TEAB prior to labelling with TMTpro 18-plex (ThermoFisher) according to manufacturer instructions. The reaction was quenched with 5% hydroxylamine (ThermoFisher) and samples combined in equal ratios as determined by running 1 μl of each sample combined on an orbitrap instrument. The SDC was precipitated from the combined samples using 2% formic acid (FA, Merck).

### Enrichment of phospho-peptides and sialylated glyco-peptides

Phospho-peptides and sialylated n-linked glyco-peptides were enriched as previously described^32^. Briefly, peptides were dissolved in titanium dioxide (TiO_2_) loading buffer (80% acetonitrile (ACN, VWR), 5% Trifluoroacetic acid (TFA, Merck), 1 M glycolic acid (Sigma)) and incubated with TiO_2_ beads (GL Sciences Inc), which bind the negatively charged phospho-peptides and sialylated glyco-peptides. The unbound “non-modified” fraction from the TiO_2_ enrichment was kept for quantitative proteomics and the modified fraction was eluted from the TiO_2_ beads using 25% ammonium hydroxide (Merck), pH 11.3, and incubated overnight with PNGase F (New England BioLabs) and sialidase A (Prozyme) for deglycosylation. The phospho-peptides were separated from the formerly sialylated glyco-peptides with a second round of TiO_2_ enrichment.

The modified peptides were purified and desalted by in-house-made reversed-phase microcolumns. Peptides were acidified and loaded onto p200 pipette tips packed with Empore SPE disks C18 (Sigma) and Oligo R3 Resin (Applied Biosystems). The non-modified peptides were similarly acidified and loaded on an Oasis HLB column (Waters), activated with methanol (VWR) and 100% ACN. The columns were equilibrated by 0.1% TFA solution. All purified peptides were eluted by 60% ACN, 0.1% TFA and dried prior to high pH fractionation.

### High pH fractionation

The 3 fractions (non-modified peptides, phospho-peptides and sialylated glycopeptides) were separately dissolved in 20 mM ammonium formate, pH 9.3, loaded on an Acquity UPLC® -Class CSHTM C18 column (Waters) and fractionated on a Dionex Ultimate 3000 HPLC system (Thermo Scientific). 20 concatenated fractions were collected for the non-modified peptides and 12 each for the phospho-peptides and sialylated glyco-peptides.

### Nano-flow liquid chromatography-mass spectrometry (nLC-MS/MS)

The samples were resuspended in 0.1% FA (buffer A) and loaded onto an in-house made two-column system containing a 3 cm pre-column (100 μm inner diameter packed with Reprosil-Pur 120 C18-AQ, 5 μm (Dr. Maisch GmbH) and an 18 cm pulled emitter analytical column (75 μm inner diameter packed with Reprosil-Pur 120 C18-AQ, 3 μm (Dr. Maisch GmbH)) on an EASY-nLC system (Thermo Scientific). The peptides were eluted with an organic solvent gradient from 100% buffer A (0.1% FA) to 40% buffer B (95% ACN, 0.1% FA) at a constant flowrate of 250 nl/min. The nLC was online connected to an Orbitrap Exploris 480 Mass Spectrometer (ThermoFisher) operated at positive ion mode with data-dependent acquisition. The Orbitrap acquired the full MS scan with an automatic gain control (AGC) target value of 3×10^6^ ions and a maximum fill time of 100ms. Each MS scan was acquired at high-resolution (120,000 full width half maximum (FWHM)) at m/z 200 in the Orbitrap with a mass range of 400-1400 Da. The 12 most abundant peptide ions were selected from the MS for higher energy collision-induced dissociation (HCD) fragmentation (collision energy: 34V). Fragmentation was performed at high resolution (60,000 FWHM) for a target of 1×10^5^ and a maximum injection time of 60 ms using an isolation window of 1.2 m/z and a dynamic exclusion. Raw data were viewed in Xcalibur v3.0 (ThermoFisher).

### Protein identification and quantification

The raw data were processed using Proteome Discoverer (v2.4, ThermoFisher) and searched against the Swissprot human database using an in-house Mascot server (v2.6, Matrix Science Ltd.). Database searches were performed with the following parameters: precursor mass tolerance of 10 ppm, fragment mass tolerance of 0.02 Da (HCD fragmentation), a maximum of 2 missed cleavages, TMT-Pro (K/N-term) and Carbamidomethyl (C) as fixed modifications, Deamidation (N) or phosphorylation (S/T/Y) as dynamic modification in the sialylated glyco-peptides and phospho-peptides, respectively. Only proteins/peptides q-value <0.01 (Percolator), Mascot rank 1 and cut-off value of Mascot score > 15 were considered for further analysis (≤1% false discovery rate (FDR)).

Only proteins with two or more unique peptides were considered for further analysis in the non-modified group. Sialylated glyco-peptides (NxS/T/C motif) were manually sorted based on information from UniProt^61^ on known glycosylation and cellular localisation (Golgi/endosome/lysosome/membrane/extracellular) to exclude spontaneous deamidations. Statistical testing to identify significant differences between the two groups was performed on all three datasets using PolyStest^62^ applying the Rank products test with FDR = 0.05 to correction for multiple testing.

Gene Ontology (GO) term enrichment analysis was performed in Cytoscape StringApp^63^ applying the ClueGO^64^ plug-in to identify enrichment of functionally grouped GO terms in the category “Biological process” on the significantly different proteins/peptides from the three datasets. For non-modified proteins and phospho-peptides fold change cut-offs of >0.3 or <- 0.3 and >0.5 or <-0.5, respectively, were applied. Enrichment was determined by two-sided hypergeometric test with Bonferroni step-down (q-value <0.05). The Homo sapiens (9606) marker set was applied with the following settings: GO term fusion selected, network confidence score 0.5, GO tree levels 6-12 and a Kappa score threshold of 0.5. For GO term selection a minimum of 3 genes and 5% of genes were used for the non-modified and sialylated glyco-proteins, whereas a minimum of 5 genes and 10% were used for the larger lists of phosphorylated proteins. Data visualisations were performed in R using ggplot2^65,66^.

### Lipid and metabolite extraction

Day 40 FOs were transferred to low-bind tubes, washed x 1 with 50 mM ammonium acetate (Sigma) and snap-frozen on dry ice. Metabolites and lipids were extracted using a modified Folch approach: FOs were sonicated 2 x 10 sec in ice-cold 1:2 methanol/chloroform solvent containing SPLASH LIPIDOMIX MS standard internal standard mix (Merck). 1:6 H_2_O was added to the samples before shaking at 1000 rpm, 4 °C, for 30 minutes, followed by centrifugation for 10 minutes at 16000 rcf, 4 °C. The metabolite-containing aqueous, upper phase and the lipid-enriched organic, lower phase were collected. The aqueous phase was re-extracted with 86:14:1 chloroform/methanol/H_2_O solvent, shaking at 1000 rpm, 4 °C, for 20 minutes, and centrifuging at 16000 rcf, 4 °C, for 10 minutes. The aqueous phase containing metabolites was collected and dried by speed vacuum centrifugation. The lower, organic phase containing remaining lipids was added to the previous organic phase and dried under a stream of nitrogen (N2) and stored at −20 °C until the day of analysis.

### Metabolomic and lipidomic analysis

Samples for metabolomics were resuspended in 0.1% formic acid 25 µl before injection of 3 µl using an Vanquish Horizon UPLC (Thermo Fisher Scientific) equipped with a analytical column(2.1 × 150 mm and 1.8 μm particle size, Agilent Technologies) operated at 40°C. The analytes were eluted using a flow rate of 400 μL/min and the following composition of eluent A (0.1% formic acid) and eluent B (0.1% formic acid, acetonitrile) solvents: 3% B from 0 to 1.5 min, 3% to 40% B from 1.5 to 3 min, 40% to 95% B from 3 to 5 min, 95% B from 5 to 7.6 min and 95% to 3% B from 7.6 to 8 min before equilibration for 3.5 min with the initial conditions. The flow from the UPLC was coupled to a TimsTOF Flex (Bruker) instrument for mass spectrometric analysis, scanning from 40-1500 mz, operated in both positive and negative ion mode using trapped ion mobility spectrometry. Collision energy of 20 and 40 Ev was applied.

Samples for lipidomics were resuspended in 25 µl chloroform/methanol (1:1) and 1 µl injected using a Vanquish Horizon UPLC (Thermo Fisher Scientific) equipped with a Waters ACQUITY Premier CSH (2.1 x 100mm, 1.7 µM) column operated at 55°C. The analytes were eluted using a flow rate of 400 μL/min. For lipids the following composition was applied of eluent A (Acetonitrile/water (60:40), 10 mM ammonium formate, 0.1% formic acid) and eluent B (Isopropanol/acetonitrile (90:10), 10 mM ammonium formate, 0.1% formic acid): 40% B from 0 to 0.5 min, 40–43% B from 0.5 to 0.7 min, 43-65% B from 0.7 to 0.8 min, 65-70% B from 0.8 to 2.3 min, 70-99% B from 2.3 to 6 min, 99% B from 6-6.8 min, 99-40% B from 6.8-7 min before equilibration for 3 min with the initial conditions.

For metabolites the following composition was applied of eluent A (0.1% formic acid in water and eluent B (Isopropanol/acetonitrile (90:10), 0.1% formic acid in acetonitrile): 3% B from 0 to 1 min, 3–40% B from 1 to 3 min, 40-95% B from 3 to 5 min, 95 % B from 5 to 7.6 min and 95 to 3% B from 7.6 to 8 min before equilibration for 3.5 min with the initial conditions.

The flow from the UPLC was coupled to a TimsTOF Flex instrument (Bruker) operated in both positive and negative ion mode using trapped ion mobility spectrometry. For lipids scanning from 100-1800 mz with a collision energy of 30/50 Ev in positive ion mode and 20/30 eV in negative ion mode. For metabolites scanning from 40-1500 mz with a collision energy of 20 and 40 Ev.

Data was processed in Metaboscape (v2023, Bruker). For lipidomics annotation was done using both an in-build rule-based annotation approach and a LipidBlast MS2 library^67^. For metabolomics annotation was done by firstly searching MS2 spectra against the following MSMS libraries: Metabobase (Bruker), National Institute of Standards and Technology 17 (NIST17) and MassBank of North America (MoNA). Next the not annotated compounds were annotated using Metfrag for in silico annotation^68^.

Features were removed if their average signal were not > 5 x more abundant in the QC samples than blanks (water extraction). The signals were normalised to internal standards in the SPLASH mix before correction for signal drift using statTarget^69^. Finally, signals were normalised using the QC samples, before log transformation (base 10) and auto scaling, all done in Metaboanalyst^70^.

### Western blotting

Day 40 and day 120 FOs were lysed in 1% SDC in 50 mM TEAB, pH8, and sonicated 2 x 10 sec on ice at 35% amplitude. Following sonication, samples were centrifuged at 5000 rcf for 10 min at RT and supernatant transferred to new Eppendorf tubes. Protein concentration was determined by Nanodrop (Implen). Proteins were separated using Bolt 4-12% Bis-Tris Plus Gels (Thermo Scientific) with samples loaded in NuPage LDS Sample buffer (Thermo Scientific) with NuPage Sample Reducing agent (Thermo Scientific) and PageRuler Plus pre-stained protein standard (Thermo Scientific) or SeeBlue™ Plus2 Pre-stained Protein Standard (Invitrogen) for size reference. Proteins were transferred to PVDF membranes using the Trans-Blot® Turbo™ Transfer System (BioRad) and membranes were blocked with 5% skimmed milk in TBST buffer for 1 h and incubated overnight at 4 °C in TBST with the following primary antibodies: Anti-rabbit TOMM20 (Abcam #ab186735) 1:1000, anti-mouse Map2a+b (Sigma #M1406) 1:500, anti-mouse ꞵ-actin HRP-linked (Abcam #ab49900) 1:50000, anti-rabbit Synapsin 1 (Abcam #ab254349) 1:1000, anti-rabbit HIF1α (Abcam #ab51608) 1:1000, anti-rabbit Glial Fibrillary Acidic Protein (GFAP, Dako #Z0334) or rabbit anti-LAMP1 (Abcam #24170) 1:500. Following 3 x wash in TBST, the membranes were incubated for 1 hr at RT in TBST with the following secondary antibodies: anti-rabbit IgG, HRP-linked (Cell signalling #7074) 1:10000 or anti-mouse IgG, HRP-linked (Abcam #ab6728) 1:10000. Following 3 x wash in TBST, the membranes were visualised with Immobilon ECL Ultra Western HRP Substrate (Millipore) using an Amersham 680 Imager (GE Healthcare). Representative full lane Western blots for all antibodies are shown in Fig. S8.

As controls for the HIF1α blots, the human ventral midbrain neural precursor (NPC) line hVM-bcl-xl was differentiated to neurons for 10 days as previously described^71^ and incubated for 4 hours at either normoxic (21% oxygen tension) or hypoxic (1%) conditions. The neurons were quickly washed once with dPBS, collected in 1% SDC in 50 mM TEAB and snap-frozen on dry ice.

### Multi-electrode array (MEA) analysis

MEA recordings were performed on a BioCam Duplex system (3Brain). The recording area of Accura HD-MEA cartridges (3Brain) were coated ON with 100 μg/ml poly-l-lysine (Sigma), washed x 3 with dPBS (ThermoFisher) and coated ON with 50 μg/ml laminin (Sigma). Day 110 FOs were plated on the HD-MEA cartridges by removing the laminin, placing the FOs gently on the recording area using a sterile spatula and removing all medium. Attachment of the FOs to the recording area was promoted by stepwise addition of small amounts of medium (10-100 μl) with 5-10 min incubation periods in between. When attachment of the FOs was ensured, the cartridge reservoir was filled with 1.5 ml medium. The FOs were cultured on the HD-MEA cartridges with medium change every 3-4 days. On day 120, 2 min recordings were performed on the BioCam Duplex system with temperature set to 37°C and spike detection in balanced mode. Analysis was performed using BrainWave 5 software averaging data from the 100 most active units from each organoid HD-MEA recording.

### Functional assays for LDH and ROS

The LDH release from day 40 FOs was measured by incubating individual FOs in 150 μl maturation medium ON and quantifying the LDH content in the medium using the LDH-Glo™ Cytotoxicity Assay (Promega) according to manufacturer instructions.

The ability of day 40 FOs to eliminate ROS was measured using the ROS-Glo® H_2_O_2_ Assay (Promega) by incubating individual FOs for 2 hours with the H_2_O_2_ substrate and subsequently detecting H_2_O_2_ levels in the medium (with and without FOs) according to manufacturer instructions.

For both assays the relative luminescence intensity was measured using a Fluostar Omega Plate reader (BMG Labtech) and normalised to the protein content of each FO as measured by Nanodrop.

### Seahorse analysis

Day 55-60 FOs were embedded in 4% (w/v) low-gelling temperature agarose in HBSS and sectioned into 150 um slices on a vibratome with the following settings: amplitude 300 μm, frequency 6 and speed 5. The slices were cultured free floating and the following day plated in Seahorse XFp Microplates (Agilent) using the same approach as described for the MEA analysis. ATP production from FO slices was measured using the Seahorse XF Real-Time ATP Rate Assay (Agilent) according to manufacturer instructions with the following modifications: the incubation time for oligomycin during the assay was increased to 15 min to allow for proper diffusion into the slices and the final toxin concentrations were increased to 5 μM for oligomycin and 2 μM for rotenone/antimycin (Agilent). The ATP production was normalised to the FO slice protein content as measured by Nanodrop (Implen).

### Single-cell RNAseq

Single-cell dissociation was performed on a total of 7-8 FOs (day 20) or 2-3 FOs (day 40) in triplicates. FOs were collected and incubated in 1 ml of Accumax (Sigma, A7089) at 37°C for 20 min with gentle agitation. At 5 min intervals, the sample tubes were flicked, and after 15 min pipetted 1 time, followed by a final pipetting of 7 times at 20 min. Clumps were allowed to settle and the supernatant passed through a 70 µm strainer (Fisherbrand, 11597522). Cells were counted (CountessII, Invitrogen) and at least 1×10^6^ viable cells were collected for the following Evercode^TM^ Cell fixation (Parse Biosciences, ECF2001) according to manufacturer instructions omitting bovine serum albumin in the pre-fixation buffer. The 12 samples were barcoded and sequencing libraries prepared using the Evercode™ WT Mini v2 kit (Parse Biosciences) according to the manufacturer’s protocol. The libraries were then sequenced on NextSeq 2000 (Illumina) using a P2 flowcell (Illumina) for 200 cycles. The Illumina sequencing generated pairs of FASTQ files. The bioinformatics pipeline split-pipe (version 1.0.6p), produced by Parse Biosciences, was used to process these files^72^. The pipeline performed multiple functions, including identifying barcodes, mapping reads to the human reference genome (GRCh38 / version 102) and quantifying gene expression at the single cell level. The pipeline was run with default parameters for a Parse Mini Kit using Version 2 chemistry.

The filtered digital gene expression (DGE) matrix was analysed in R using Seurat^65,66,73^. Only cells with total transcripts >100 and <150,000, unique transcripts <12,000 and mitochondrial transcripts <10% of total were analysed (Fig. S5A). The full dataset was split into the four conditions, normalising and identifying variable features of each dataset before reintegration based on repeatedly variable features (integration anchors). The integrated dataset was scaled and dimensionality reduction performed using principal component analysis (PCA) and visualised with UMAP generated on the first 15 principle components (Fig. S5B-C). K-Nearest Neighbour graph-based clustering, with Louvain algorithm and resolution = 1, resulted in 21 clusters, which upon differential gene expression analysis of known marker genes and literature review were combined and manually annotated to 12 clusters.

Pseudotime analysis was performed using Monocle3 on the data from guided and unguided FOs separately by subsetting and converting the integrated Seurat object^74,73^. UMAP coordinates and clusters were assigned from Seurat and trajectory analysis performed not considering cluster partitions. Cells were ordered by denoting clusters of mitotic radial glia as starting points.

### Statistical analyses

Data visualisation and statistical testing was performed in Prism (v10.0, GraphPad Software) unless otherwise stated in the above. Analysis was performed in GraphPad Prism version 5.0 (GraphPad Software) using two-tailed paired Student’s t-tests with Benjamini-Hochberg correction for multiple testing where appropriate. Results are expressed as mean ± SEM, and p- or q-values ≤ 0,05 were considered statistically significant. Statistical details for each experiment can be found in the figure legend.

## Materials

IMR90 clone 4 (WiCell #iPS(IMR90)-4)

Matrigel® Growth factor reduced (GFR) (Corning #356230)

mTESR1 medium (Stem Cell Technologies #85870)

Gentle cell dissociation reagent (Stem Cell Technologies #07174)

Cerebral Organoid Kit (Stem Cell Technologies #08570)

Y-27632, ROCK inhibitor (Stem Cell Technologies #72302)

Aggrewell^TM^800 24-well plates (Stem Cell Technologies **#**34815)

Anti-Adherence Rinsing Solution (Stem Cell Technologies #07010)

Matrigel® (Corning #356234)

Penicillin/Streptomycin (Gibco #15400)

Amphotericin B (Thermo Scientific #15290026)

STEMdiff Dorsal FO Differentiation Kit (Stem Cell Technologies #08620)

STEMdiff Neural Organoid Maintenance Kit (Stem Cell Technologies #100-0120)

Distilled Phosphate Buffered Saline (dPBS, ThermoFisher #14190)

Triethylammonium bicarbonate (Sigma #T7408)

Sodium deoxycholate (SDS, Sigma #D6750)

Dithiothreitol (DTT, Sigma #D9163)

Iodoacetamide (Sigma #I1149)

NanoPhotometer N60/N50 (Implen)

Methylated trypsin (made in-house)^75^

Acetonitrile (VWR #83640.320)

Trifluoroacetic acid (Merck #1.08178.0050)

TMTpro 16plex (Thermo #A44520)

TMTpro-134C & TMTpro-135N Label Reagents (Thermo #A52046)

Hydroxylamine (Thermo #90115)

Formic acid (Merck #1.11670.0250)

Glycolic acid (Sigma #124737)

Titanium dioxide beads, 5 µm (GL Science #5020-75010)

PNGaseF (New England BioLabs #P0705L)

Sialidase A (Prozyme #GK80040)

Ammonium hydroxide solution (Merck #1.05428.0500)

Oasis HLB column (Waters #186000132)

Methanol (VWR #20864.320)

Empore^TM^ SPE disks C8 (Sigma #66882-U)

Empore^TM^ SPE disks C18 (Sigma #66883-U)

Oligo^TM^ R3 material (Applied Biosystems #1-1339-03)

Ammonium acetate (Sigma #73594)

Acquity UPLC® -Class CSHTM C18 column (Waters)

Dionex Ultimate 3000 HPLC system (Thermo Scientific)

SPLASH LIPIDOMIX MS standard internal standard mix (Merck #330707-1EA)

Bolt 4-12% Bis-Tris Plus Gels (Thermo Scientific #NW04125BOX)

NuPage LDS Sample buffer (Thermo Scientific # #NP0007)

NuPage Sample Reducing agent (Thermo Scientific # #NP0004)

PageRuler Plus pre-stained protein ladder (Thermo Scientific #26620)

SeeBlue™ Plus2 Pre-stained Protein Standard (Invitrogen #LC5925)

Immobilon ECL Ultra Western HRP Substrate (Millipore #WBULS0100)

Trans-Blot® Turbo™ Transfer System (BioRad #1704150)

CCD camera, Amersham 680 Imager (GE Healthcare #29270769)

Trizma base (Sigma #93352)

LDH-Glo™ Cytotoxicity Assay (Promega #J2380)

ROS-Glo® H2O2 Assay (Promega #G8820)

Fluostar Omega Plate reader (BMG Labtech)

Agarose, low-gelling temperature (Sigma #A9414)

Hanks’ Balanced Salt Solution (HBSS, ThermoFisher #14025)

Leica VT1000S Vibratome (Leica Biosystems)

Seahorse XF Real-Time ATP Rate Assay Starter Pack (Agilent #103677-100)

Seahorse XFe96 FluxPak (Agilent #102601-100)

Ammonium acetate (Sigma #73594)

BioCam Duplex MEA system (3Brain)

Accura HD-MEA cartridges (3Brain)

BrainWave 5 software (3Brain)

Poly-l-lysine hydrobromide (PLL, Sigma #P1274)

Laminin (Sigma #L2020)

Formaldehyde Solution 16% w/v (Thermo Scientific #28908)

Phosphate buffered saline (Gibco #70011-036)

OCT Mounting media (VWR, #361603E)

Leica CM1860 cryostat (Leica Biosystems)

Superfrost® Plus Microscope slides (Thermo Fisher Scientific, #J1800AMNZ)

TritonX100 (Plusone #17-1315-01)

Donkey serum (BioWest, #S2170-100)

Goat serum (Gibco, #16210064)

ProLong Diamond Antifade mountant (Invitrogen #P36965)

