## Supplemental Figures for "Multi-omic analysis of guided and unguided forebrain organoids reveal differences in cellular composition and metabolic profiles"

### Supplementary information

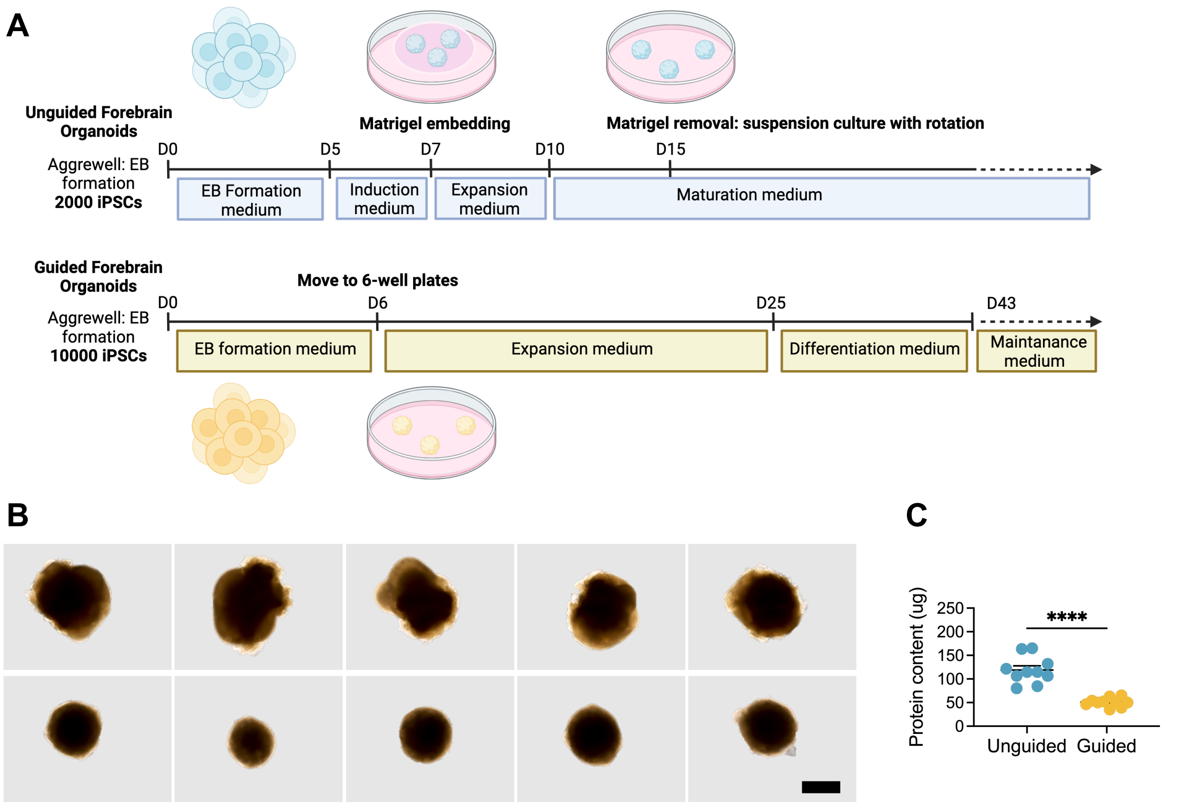


**Figure S1: overview of guided and unguided forebrain organoid (FO) differentiation and resulting size differences.**

(A) Overview of guided and unguided FO differentiation protocols based on Stem Cell Technologies kits.

(B) Representative bright field images of day 40 guided and unguided FOs. Scalebar = 1 mm.

(C) Protein content (µg) in guided and unguided FOs from two independent differentiations. Mean ± SEM, ****p≤0.0001 (n=10, Student's T-test).

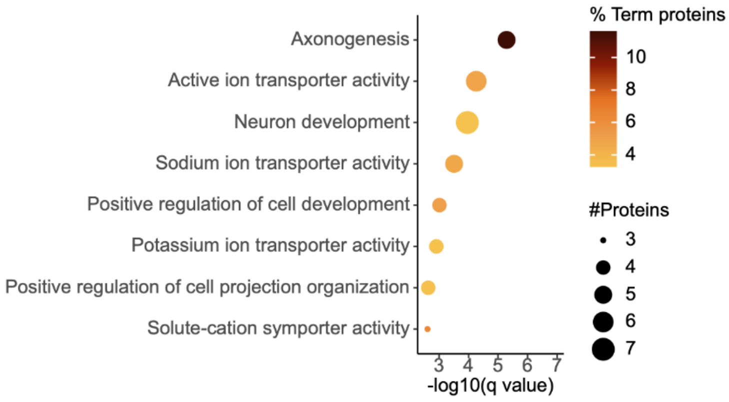


**Figure S2: Axogenesis and neuron development enrichment amongst proteins with increased sialylation levels in guided forebrain organoid (FOs).**
GO term enrichment analysis listing the pathways that were significantly enriched ($q \leq 0.05$), amongst the proteins with increased sialylation levels in guided vs unguided FOs with the dot size signifying the number of significantly different proteins in the pathway and the colour indicating how many percent these constitute out of the total number of proteins in the pathway (two-sided hypergeometric test with Bonferroni step-down).

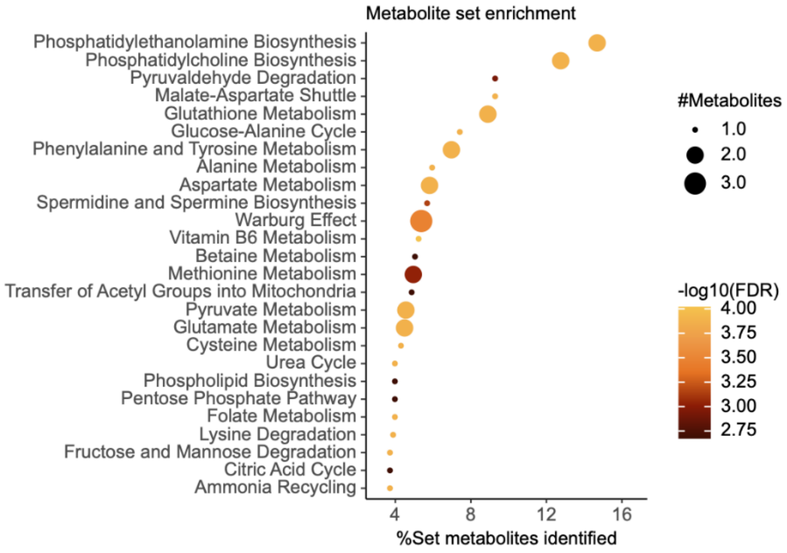


**Figure S3: The metabolomic data was enriched in metabolites related to phospholipid biosynthesis, amino acid metabolism and glucose metabolism.**
GO term enrichment analysis of all metabolites identified and annotated in guided and unguided forebrain organoid (FOs, n=5) with the dot size signifying the identified number of metabolites of each metabolite set, how many percent these constitute out of the total metabolite number of the set and colour indicating the strength of the enrichment (-log10(FDR), two-sided hypergeometric test with Bonferroni step-down).

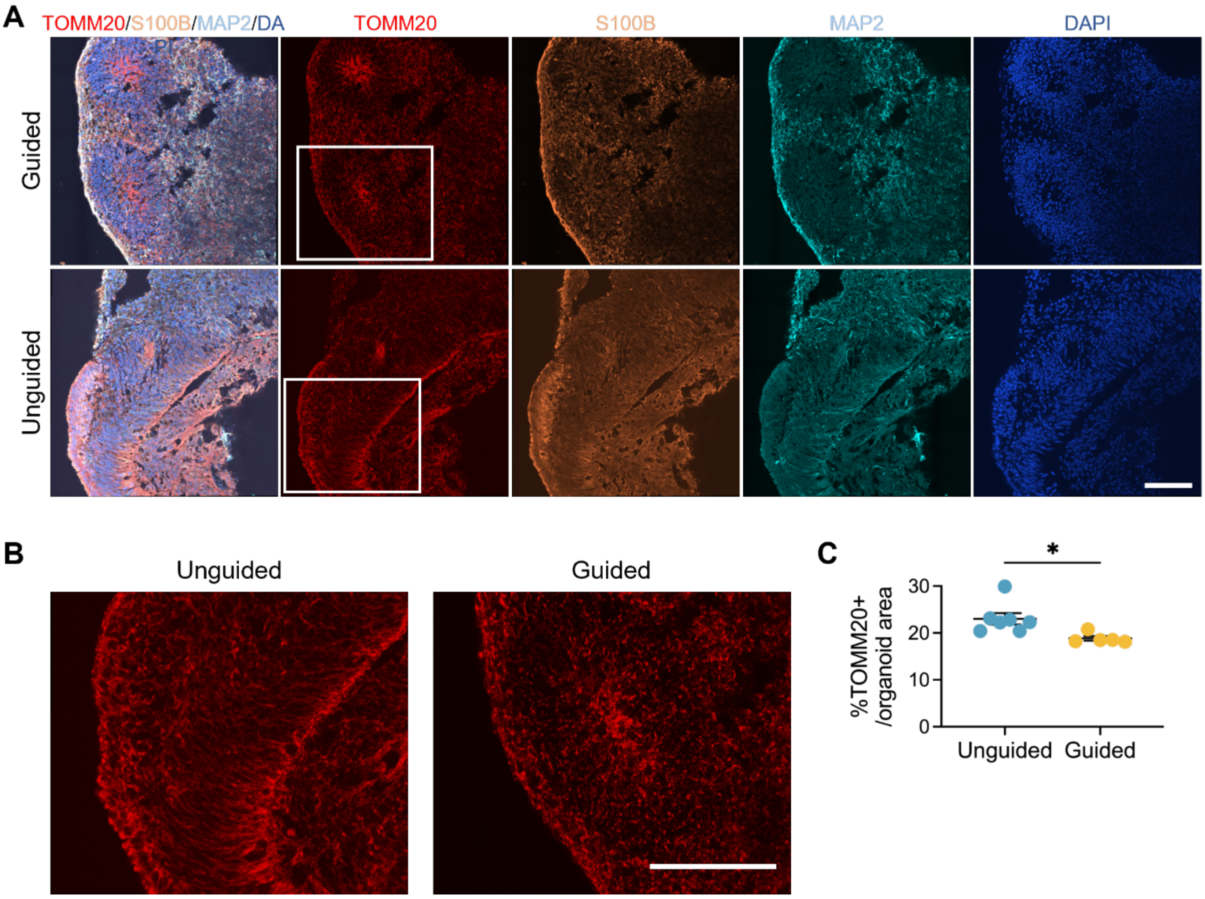


**Figure S4: Increased TOMM20 expression in unguided forebrain organoid (FOs).**
(A) ICC labelling of day 40 guided and unguided FOs for TOMM20 (red), S100B (orange), MAP2 (cyan) and DAPI (dark blue). Scalebar = 100 µm.
(B) Cut-out of TOMM20 in higher magnification. Scalebar = 100 µm.
(C) Quantification of TOMM20+ particles as % of total organoid area. Mean ± SEM, *p≤0.05 (n = 5-7 images from 2 guided/unguided FOs, Student's T-test).

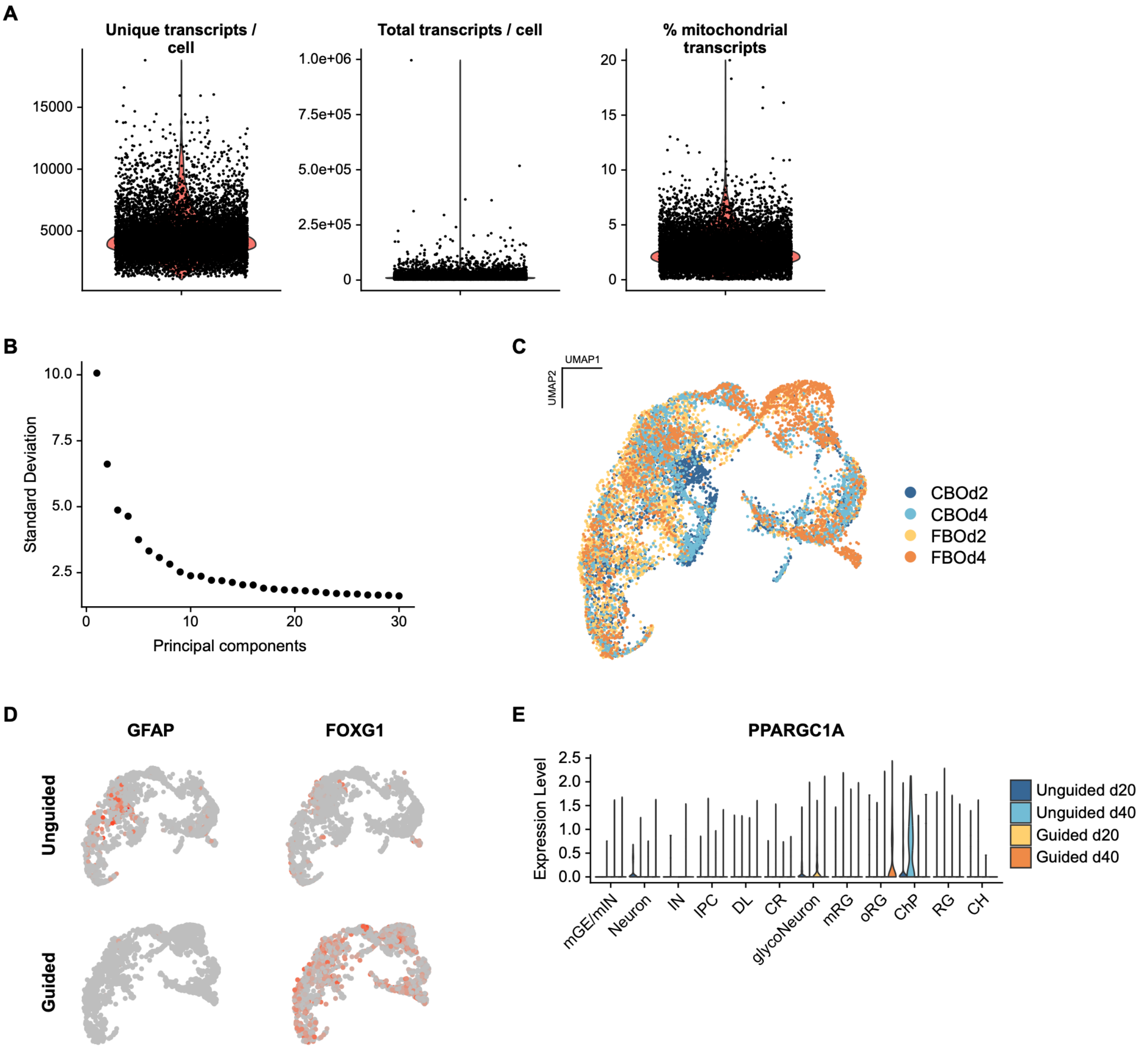


**Figure S5: Single cell transcriptomics (scRNAseq) on day 20 and 40 guided and unguided forebrain organoid (FOs)**

(A) Violin plots of numbers of unique transcripts per cell, total number of transcripts per cell and the percentage of mitochondrial transcripts out of total transcripts per cell. Based on these cut-offs of c total transcripts >100 and <150,000, unique transcripts <12,000 and mitochondrial transcripts <10% of total were chosen.

(B) Elbow plot of the standard deviation resulting from each principal component. Based on this the first 15 principal components were utilised for the UMAP.

(C) UMAP of the scRNAseq data analysed by splitting the dataset into the four conditions, normalising and identifying variable features of each dataset before reintegrating based on repeatedly variable features, showing the sample identity of each cell.

(D) Feature plots showing expression levels of GFAP and FOXG1 in day 40 unguided and guided FOs.

(E) Violin plot of RNA expression levels of PPARGC1A across the different cell types. Cell type clusters: CR; Cajal Retzius cells, RG; radial glia, oRG; outer RG, mRG; mitotic RG; CH; cortical hem, ChP; choroid plexus, DL; deep layer neurons, IPC; intermediate progenitors, IN; interneurons, mGE/mIN; medial ganglionic eminence/migratory INs.

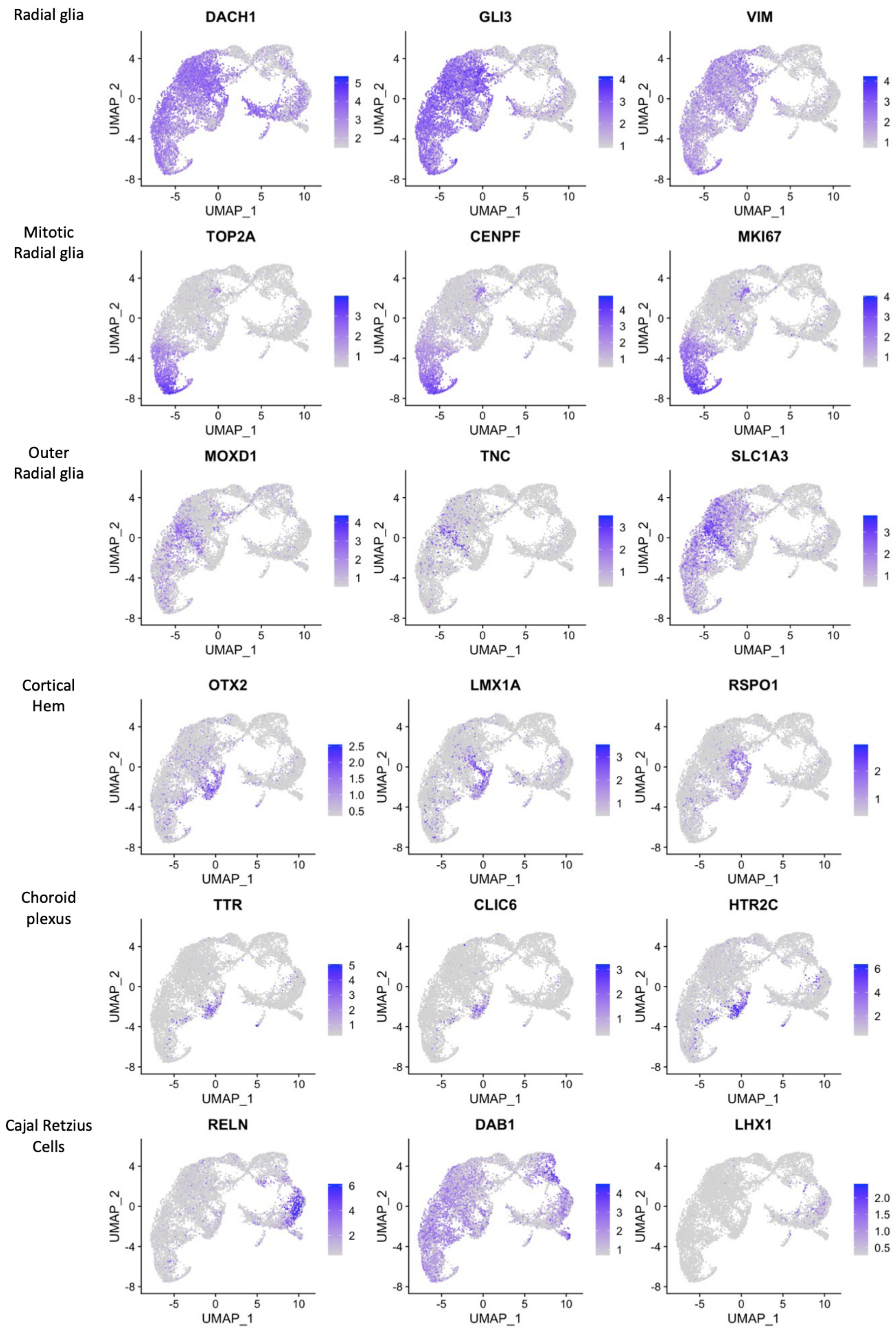

Radial glia
DACH1
GLI3
VIM
Mitotic Radial glia
TOP2A
CENPF
MKI67
Outer Radial glia
MOXD1
TNC
SLC1A3
Cortical Hem
OTX2
LMX1A
RSPO1
Choroid plexus
TTR
CLIC6
HTR2C
Cajal Retzius Cells
RELN
DAB1
LHX1
UMAP_1
UMAP_2

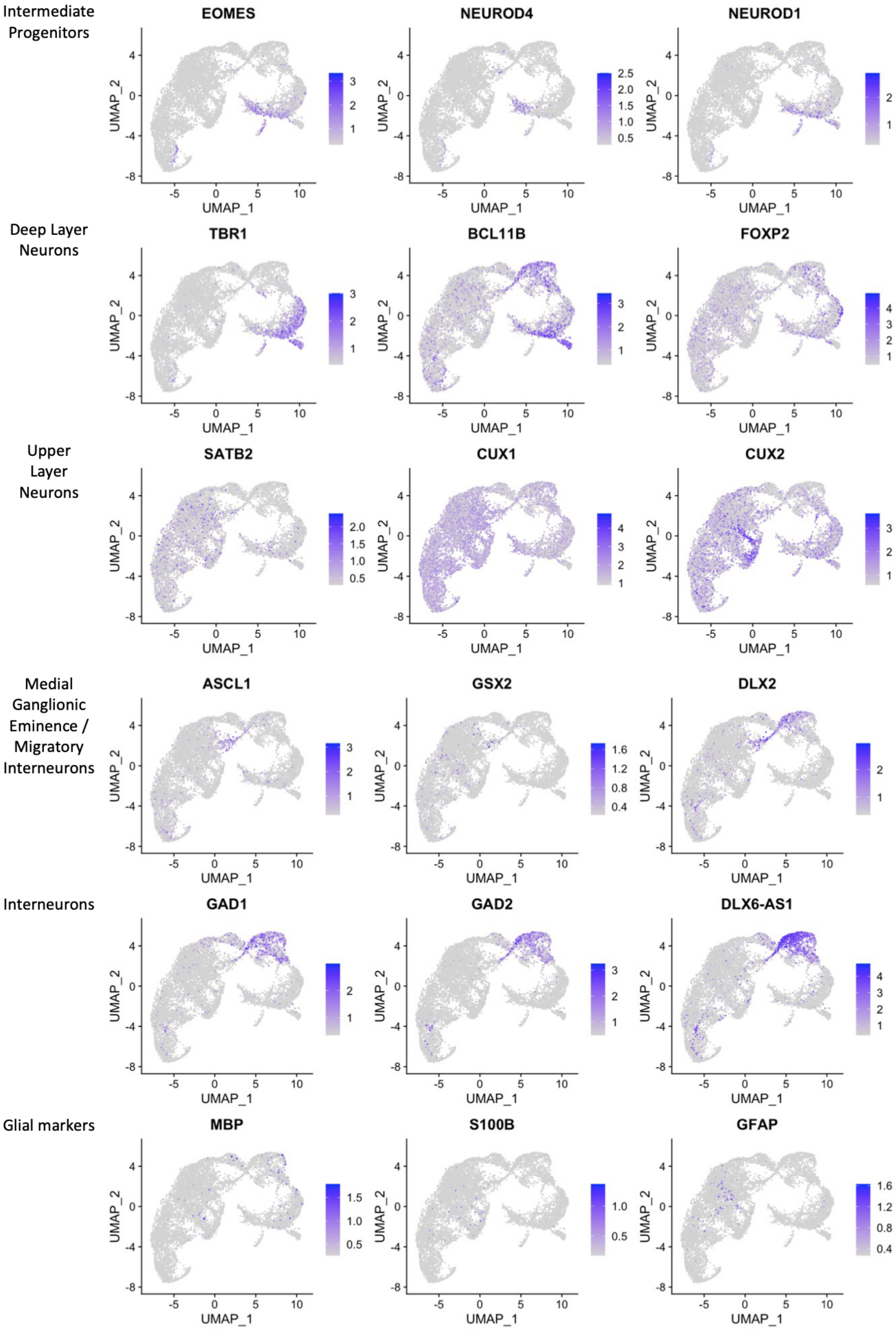
Intermediate Progenitors
EOMES
NEUROD4
NEUROD1
Deep Layer Neurons
TBR1
BCL11B
FOXP2
Upper Layer Neurons
SATB2
CUX1
CUX2
Medial Ganglionic Eminence / Migratory Interneurons
ASCL1
GSX2
DLX2
Interneurons
GAD1
GAD2
DLX6-AS1
Glial markers
MBP
S100B
GFAP
UMAP_1
UMAP_2

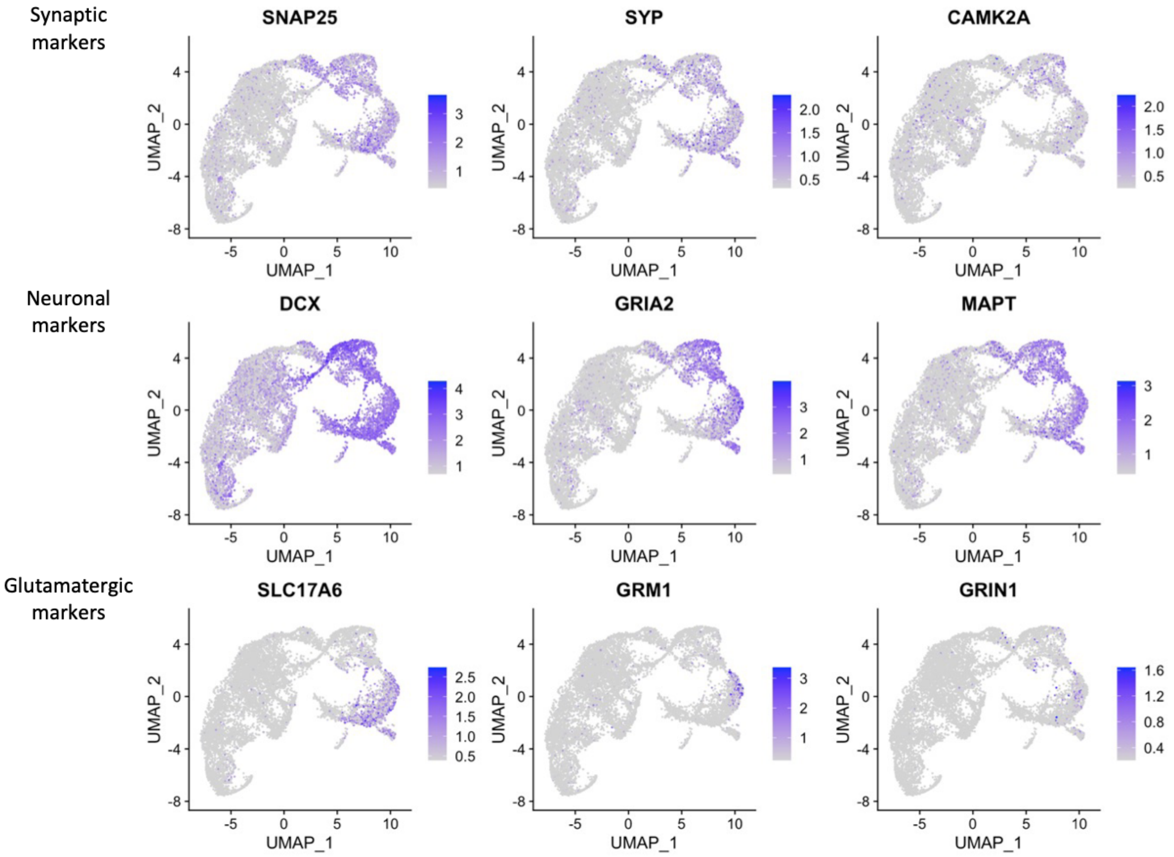


**Figure S6: Feature plots of RNA expression levels for selected markers from single cell RNA sequencing on day 20 and 40 guided and unguided forebrain organoid (FOs)**

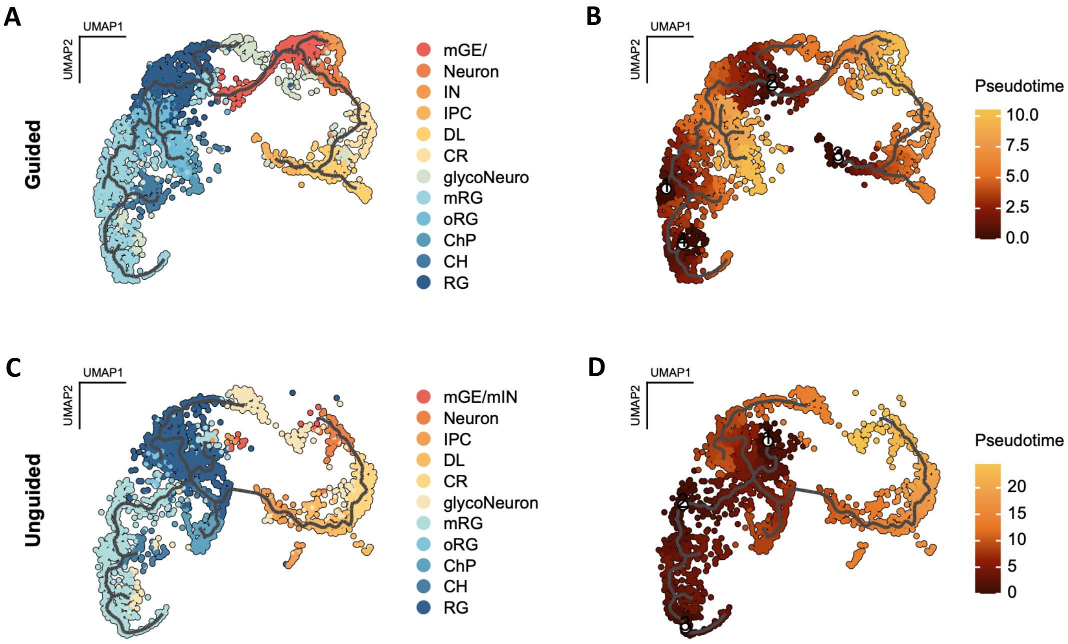


**Figure S7: Pseudotime analysis on single cell transcriptomics data from day 20 and 40 guided and unguided forebrain organoid (FOs)**

(A-D) Pseudotime analysis on guided (D-E) and unguided (F-G) FOs scRNAseq showing the developmental trajectories between the cell clusters when using mitotic radial glia, and for guided FOs also intermediate progenitors, as starting points (D,F) and the resulting pseudotime trajectories (E,G). Cell type clusters: CR; Cajal Retzius cells, RG; radial glia, oRG; outer RG, mRG; mitotic RG; CH; cortical hem, ChP; choroid plexus, DL; deep layer neurons, IPC; intermediate progenitors, IN; interneurons, mGE/mIN; medial ganglionic eminence/migratory INs.

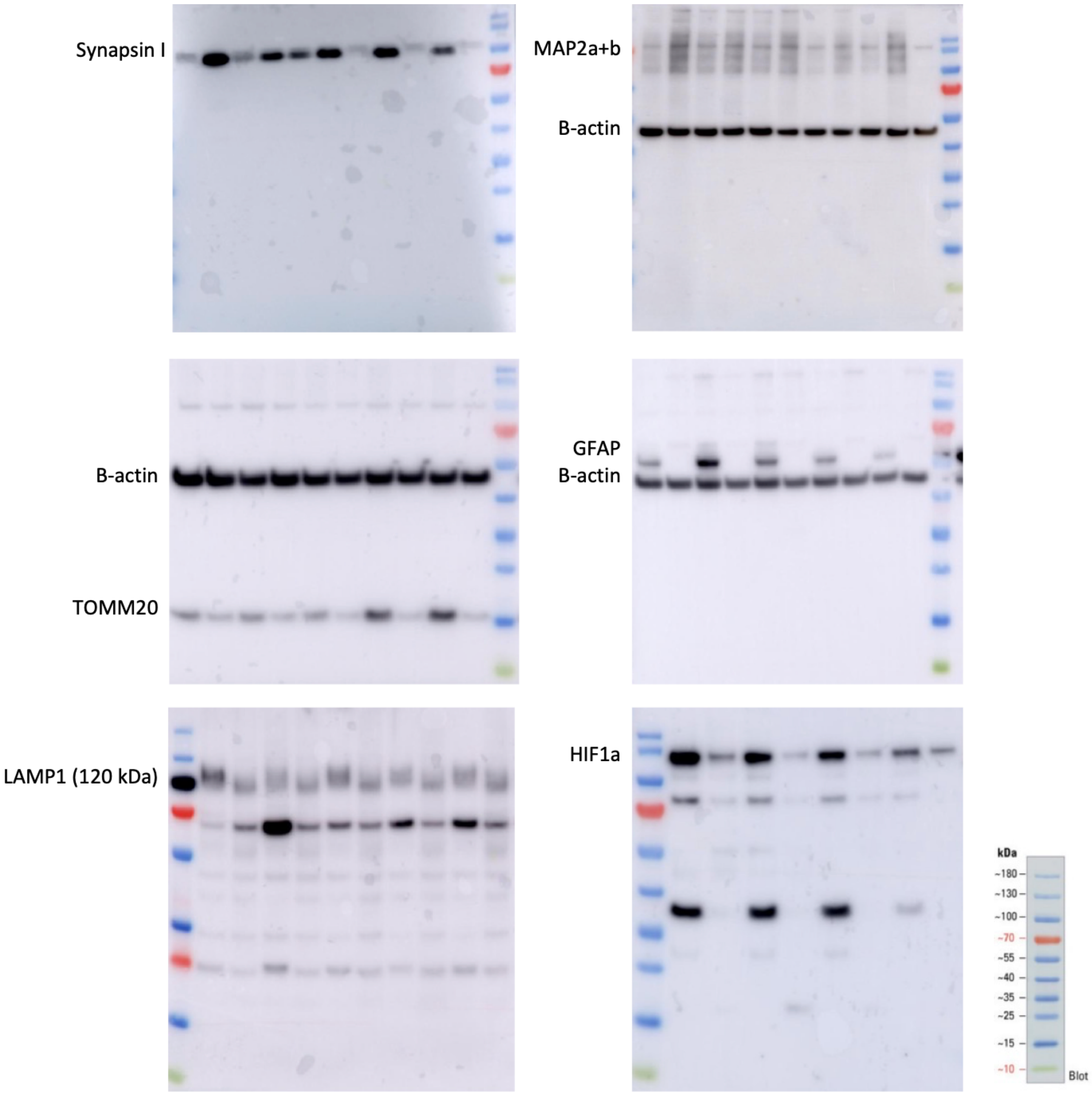


**Figure S8: Representative full lane Western blots indicating the kDa size of the quantified bands.**

**Supplementary tables**

**Table S1: Proteomic analysis on day 40 guided and unguided forebrain organoid (FOs) from 3 independent differentiations (n = 9 per group)**
(A) All non-modified proteins identified with one or more unique peptides.
(B) Non-modified proteins with significantly different abundance levels as determined by Rank products test (FDR < 0.05).
(C) All phospho-peptides with modification site specified.
(D) Phospho-with significantly different abundance levels as determined by Rank products test (FDR < 0.05).
(E) All sialylated N-linked glyco-peptides with correct motif (NxS/T/C) and relevant cellular localisation.
(F) Sialylated N-linked glyco-peptides with significantly different abundance levels as determined by Rank products test (FDR < 0.05).

**Table S2: Metaboliomics on day 40 guided and unguided FOs from one differentiation (n = 5 per group)**
(A) All identified metabolites.
(B) All annotated metabolites.

**Table S3: Lipidomics on day 40 guided and unguided FOs from one differentiation (n = 5 per group)**
(A) All identified lipids.
(B) All annotated lipids.
(C) Annotated lipids sorted according to lipid classes.

**Table S4: Differentially expressed genes for the 12 clusters identified from single cell RNA sequencing on day 20 and day 40 guided and unguided FOs from one differentiation (n = 3 per group)**
CR; Cajal Retzius cells, RG; radial glia, oRG; outer RG, mRG; mitotic RG; CH; cortical hem, ChP; choroid plexus, DL; deep layer neurons, IPC; intermediate progenitors, IN; interneurons, mGE/mIN; medial ganglionic eminence/migratory INs.
